# *Sinorhizobium meliloti* possesses a complete Embden-Meyerhoff-Parnas pathway that is indispensable for symbiotic nitrogen fixation

**DOI:** 10.64898/2026.05.26.727922

**Authors:** Ryan D. J. Payton, Sabhjeet Kaur, George C. diCenzo, Ivan J. Oresnik

## Abstract

Rhizobia are an agronomically valuable group of bacteria capable of entering into endosymbiotic relationships with leguminous plants, during which they fix atmospheric nitrogen using energy derived from the metabolism of plant-provided dicarboxylic acids. It is generally assumed that the gluconeogenic catabolism of dicarboxylic acids proceeds via the Embden-Meyerhoff-Parnas pathway in rhizobia. However, rhizobia are classically thought to lack the phosphofructokinase enzyme required for conversion of fructose-1,6-bisphosphate to fructore-6-phosphate as part of this pathway. Here, we demonstrate that a model rhizobium, *Sinorhizobium meliloti*, encodes a phosphofructokinase, completing the Embden-Meyerhoff-Parnas pathway of this organism. Biochemical characterization of the *S. meliloti* phosphofructokinase demonstrates that it can catalyze the reversible phosphorylation of fructose-6-phosphate under *in vitro* conditions in a pyrophosphate-dependent, rather than ATP-dependent, manner. We further show that *S. meliloti* also encodes a distinct fructose-1,6-bisphosphatase that can phenotypically complement the loss of the phosphofructokinase enzyme. Loss of both enzymes results in a block of the gluconeogenic pathway in *S. meliloti* and results in *S. meliloti* being unable to fix nitrogen in symbiosis with alfalfa (*Medicago sativa*). Phylogenetic analyses and complementation studies demonstrate that PPi-dependent phosphofructokinases are broadly distributed across the phylum *Pseudomonadota* (syn. *Proteobacteria*), including most rhizobial species of the class *Alphaproteobacteria*, suggesting both that PPi-dependent phosphofructokinases are likely more broadly distributed than is generally recognized, and that the catabolism of dicarboxylic acids in most rhizobia proceeds via a PPi-dependent phosphofructokinase.

**SIGNIFIGANCE:** Central carbon metabolism is an important biochemical network that bridges the gap between substrate catabolism and biosynthetic reactions in all living organisms. However, much of what we know about metabolism comes from the study of a few model organisms such as the bacterium *Escherichia coli*. Here, we identified the enzyme catalyzing a key step of central carbon metabolism in rhizobia (nitrogen-fixing bacterial symbionts of legumes), which until now had remained undetected. We show that this enzyme is dependent on pyrophosphate, which is different than the situation in *E. coli*, helping to explain why previous studies failed to identify this enzyme in rhizobia and highlighting the limitations associated with generalizing our understanding of metabolism from a limited subset of organisms.

## INTRODUCTION

Nitrogen is a growth-limiting nutrient for plants and is essential in agriculture for maintaining crop yields. Biologically usable nitrogen can be produced either through biological nitrogen fixation or via the industrial Haber-Bosch process (1). It is estimated that the use of industrially produced nitrogen in agriculture has increased the global population, such that the nutritional requirements of approximately 50% of the current population is met as a direct consequence of this practice (2–4). However, the manufacturing and application of synthetic fertilizers are energy-intensive, generate greenhouse gases, and damage ecosystems (3, 5, 6).

Symbiotic nitrogen fixation (SNF) is a process in which specific soil microbes (e.g., rhizobia) associate with specific plants (e.g., legumes) and convert N_2_ into a bioavailable form (ammonia), which is then supplied to the plant hosts in exchange for reduced carbon (7). Under ideal circumstances, SNF can replace the need for nitrogen fertilization of economically important legume crops such as soybean (*Glycine max*) and peas (*Pisum sativum*), thereby lowering greenhouse gas emissions and promoting sustainability in the cultivation of these crops. Consequently, there is significant scientific interest in fully understanding the molecular and metabolic mechanisms that underpin SNF to engineer improved nitrogen-fixing symbioses in legume crops and to transfer SNF to staple non-legume crops like cereals (8).

As SNF is built on a metabolic exchange between the plant and bacterial partners, understanding the metabolic pathways that are active during SNF is core to being able to engineer this process. A commonly used model system for studying metabolism during SNF consists of the rhizobium *Sinorhizobium meliloti* strain Rm1021 and its host legumes *Medicago sativa* (alfalfa) *and M. truncatula* (barrel medic) (9–11). The central carbon metabolic pathways of *S. meliloti* Rm1021 include the Embden-Meyerhof-Parnas (EMP) pathway, the Entner-Doudoroff (ED) pathway, the pentose phosphate (PP) pathway, and the tricarboxylic acid (TCA) cycle (**Figure 1**) (10). Past studies using ^13^C labeling experiments have shown that glycolytic growth occurs via the ED, PP, and TCA pathways, while the upper part of the EMP pathway is exclusively used for gluconeogenesis (12–14)

**Figure 1.**
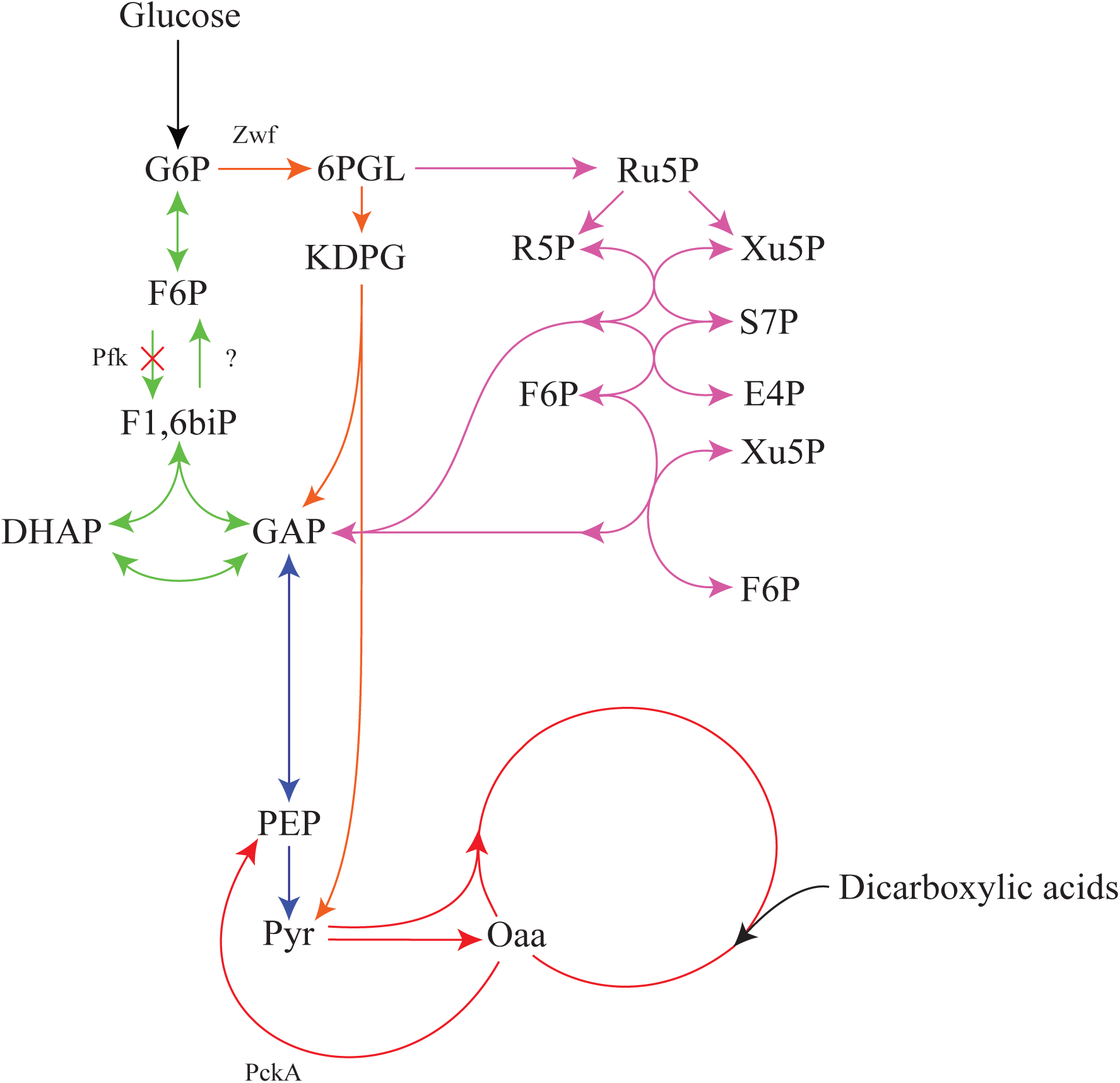
Reaction pathways of central carbon metabolism in *Sinorhizobium meliloti*. Arrows indicate reaction directionality, and reactions of interest are labelled with the proteins capable of catalyzing the reaction. Double sided arrows represent reversible reactions, while single sided arrows are irreversible. Major pathways are indicated by arrow colour: Upper Embden-Meyerhof- Parnas pathway, green; Lower Embden-Meyerhof-Parnas pathway, blue; Entner-Doudoroff pathway, orange; Pentose phosphate pathway, pink; Tricarboxylic acid cyle, red. Peripheral reactions and other metabolic bypasses are colored black. Metabolites are abbreviated as follows: Glucose-6-phosphate (G6P); Fructose-6-phosphate (F6P); Fructose-1,6-bisphophate (F1,6biP); Dihydroxyacetone phosphate (DHAP); Glyceraldehyde-3-phosphate (GAP); 6- phosphoglucolactone (6PGL); 6-phosphogluconate (6PG); Ribulose-5-phosphate (Ru5P); Ribose- 5-phosphate (R5P); Xylulose-5-phosphate (X5P); Seduheptulose-7-phosphate (S7P); Erythrose-4- phosphate (E4P).

A conserved feature of rhizobium-legume symbioses is that the nitrogen-fixing bacteria receive dicarboxylic acids (primarily malic acid) as the primary source of carbon from the legumes (15, 16). Catabolism of these dicarboxylic acids is essential to support the symbiosis, as mutation of genes encoding the dicarboxylic acid transporter results in a complete loss of nitrogen fixation (16). It is believed that carbon flow from the dicarboxylic acids is limited to the TCA cycle and the lower EMP pathway in nitrogen fixing rhizobia (17). Indeed, mutations in genes encoding enzymes for the lower EMP pathway, as well in the TCA cycle have been shown to form ineffective nodules that do not fix nitrogen (18–21). Interestingly, *S. meliloti* strains carrying mutations in genes encoding enzymes of the upper EMP pathway or the PP pathway, including phosphoenolpyruvate carboxykinase (*pck*), triose-phosphate isomerase and tetrulose phosphate isomerase (*tpiA, eryH*), or suppressors of a transketolase (*tktA*), all form functional nodules but have reduced efficiency of nitrogen fixation (18, 22, 23). Collectively, these observations suggest that the upper EMP pathway and PP pathway also affect nitrogen fixation in ways that remain poorly understood.

The enzyme phosphofructokinase (PFK) catalyzes the conversion of fructose-6-phosphate to fructore-1,6-bisphosphate as part of the EMP pathway. This enzyme links the upper and lower halves of the EMP pathway and is generally essential for the functioning of the EMP pathway (24). In addition, its allosteric regulation serves as a critical mechanism to control the flow of carbon through point in the EMP pathway (25). However, *S. meliloti* and most other rhizobia lack open reading frames encoding an ATP-dependent phosphofructokinase (*pfk*), and its activity has never been detected through functional assays despite multiple attempts (10, 26–28). In addition, a fructose-1,6-bisphosphatase (*fbp*) that catalyzes the conversion of fructose-1,6-bisphosphate [F1,6biP] to fructose-6-phosphate [F6P] in the EMP pathway has been assayed and its activity detected, yet the gene encoding this activity has never been identified (10, 18). Considering the importance of dicarboxylic acid metabolism to rhizobia and SNF, and that the EMP pathway is essential for gluconeogenesis and growth with dicarboxylic acids, not understanding how rhizobia convert fructose-1,6-bisphosphate to fructore-6-phosphate during gluconeogenesis represents a major knowledge gap.

In this work, we demonstrate that the conversion of fructose-1,6-bisphosphate to fructore- 6-phosphate occurs primarily via a pyrophosphate-dependent phosphofructokinase (Pfp) in *S. meliloti* and other rhizobia. We further show that in *S. meliloti*, the loss of Pfp activity can be complemented by a specific fructose-1,6-bisphosphatase (Fbp). We additionally demonstrate that the ability of *S. meliloti* to convert fructose-1,6-bisphosphate to fructore-6-phosphate is essential for SNF as deletion of both *pfp* and *fbp* result in a non-effective symbiosis. These results allowed us to fill in the last major gap in the central carbon metabolic pathways of *S. meliloti* and other rhizobia, and gain new insight into the metabolic processes supporting SNF by rhizobia.

## RESULTS

### *S. meliloti* encodes a pyrophosphate-dependent phosphofructokinase

The *S. meliloti* Rm1021 genome lacks an open reading frame predicted to encode an ATP-dependent phosphofructokinase (*pfk*). On the other hand, the original *S. meliloti* genome annotation suggested that *smc01852* may encode a pyrophosphate-dependent phosphofructokinase (*pfp*) (26). Likewise, a comprehensive *S. meliloti* genome-scale metabolic model also suggested that *smc01852* encodes a Pfp, but the model excludes *smc01852* from the core metabolic model and notes that this annotation lacks experimental support (29). Lastly, querying the insertion-sequencing (INseq) results summarized in the Fitness Browser (30) suggested that mutation of *smc01852* negatively impacts growth on multiple carbon sources, with the most severe phenotypes on glycerol, L-rhamnose, m-inositol, D-galactose, and D,L-lactate. Collectively, these preliminary findings suggested that *smc01852* may encode a Pfp involved in carbon metabolism.

To experimentally test whether *smc01852* encodes a functional pyrophosphate-dependent phosphofructokinase, the corresponding product was purified for *in vitro* assays (**Appendix S1, Figure S1**). The purified protein could catalyze both the phosphorylation of F6P (glycolytic direction) and dephosphorylation of F1,6biP (gluconeogenic direction) (**Figure 2**), with the turnover efficiencies suggesting that Pfp is poised in a glycolytic direction in the absence of allosteric regulation (**Figure 2**). We were unable to determine the affinity of the enzyme for orthophosphate (Pi) using our conditions because of contaminating Pi from the elution buffer that carried over during the purification. Additionally, it was found that ATP could not replace PP_i_ as a phosphate donor for F6P. Together, these findings show that *smc01852* encodes a bidirectional PP_i_-dependent phosphofructokinase.

**Figure 2.**
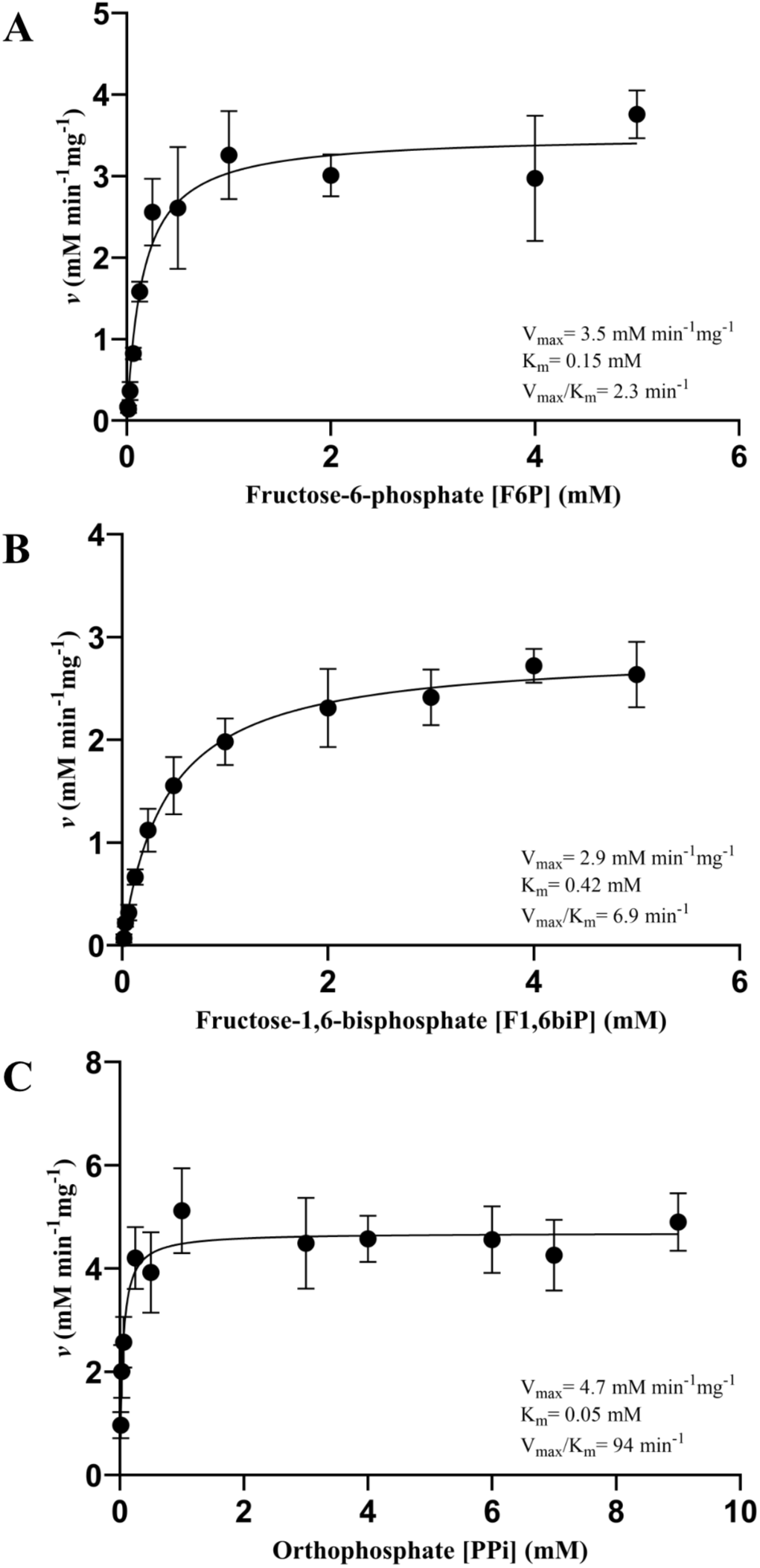
The *Sinorhizobium meliloti* Pfp enzyme functions as a phosphofructokinase *in vitro*. Data generated for the (**A**) phosphorylation of fructose-6-phosphate (F6P), (**B**) dephosphorylation of fructose-1,6-bisphosphate (F1,6biP), or (**C**) affinity for orthophosphate (PPi) were fitted to Michalis-Menten models. Dots represent the mean of technical triplicates, error bars denote standard deviations, and curves show the fitted equation. K_m_ and V_max_ were determined by linear regression analysis using *Graphpad Prism* version 9.3.1, and they represent the average of at least three independent experiments. Plots were generated using *Graphpad prism* 9.3.1.

### Pfp is a non-essential enzyme involved in the EMP pathway of *S. meliloti*

We generated unmarked (Δ*pfp*) and marked (Δ*pfp*::Nm^R^) deletions of *smc01852* (*pfp*) to test whether Pfp is involved in the EMP pathway. Unexpectedly, our results demonstrated that Pfp is not required for a functional EMP pathway in *S. meliloti*, as both mutants displayed wildtype growth on defined medium supplemented with various carbon sources, including glucose, succinate, L-arabinose, fructose, glycerol, ribose, xylose, and erythritol. To further probe whether Pfp is involved in the EMP pathway in a non-essential capacity, we performed targeted metabolomics to identify whether deletion of *pfp* resulted in altered metabolic pools. Deletion of *pfp* resulted in an accumulation of multiple intermediates of central carbon metabolism (**Appendix S1, Figure S2 and Table S1; Dataset S1**). Notably, F1,6biP, which is a substrate for Pfp when functioning in the gluconeogenic direction, was found to be ∼ 10-fold higher in the mutant than in the wild type when grown in either glucose or succinate (**Figure 3**), suggesting that the accumulation is directly associated with the mutation and providing evidence that Pfp is involved in gluconeogenesis and the EMP pathway in *S. meliloti*.

**Figure 3.**
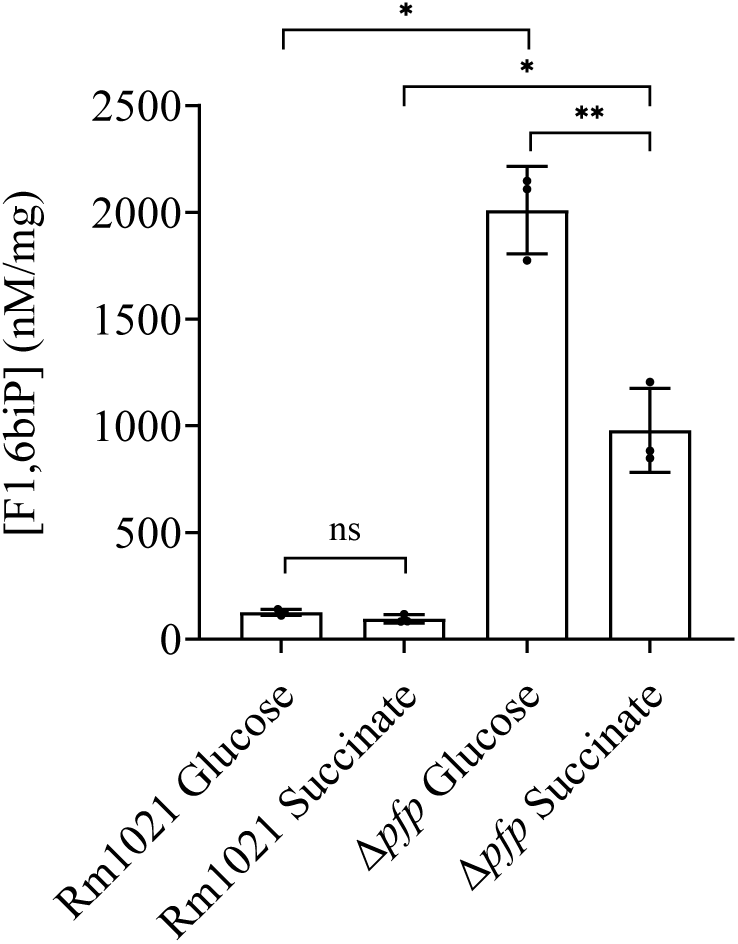
Pfp contributes to *in vivo* fructose-1,6-bisphosphate metabolism in *Sinorhizobium meliloti*. The concentrations of fructose-1,6-bisphosphate in wildtype *S. meliloti* Rm1021 and a *S. meliloti* Δ*pfp* mutant (strain SRmD674) when grown with glucose or succinate as the sole carbon source are shown. * Denotes a statistically significant difference (P < 0.05) between different strains determined using Student’s t-test. ** Indicates concentrations of fructose-1,6-bisphosphate that are significantly different in the same strain grown with glucose or succinate. “ns” corresponds to no statistical difference between strains.

### A fructose-1,6-bisphosphatase compensates for the loss of Pfp in *S. meliloti*

It is well documented that *S. meliloti* exhibits genetic redundancy in core metabolic pathways, which can mask phenotypes unless all involved genes are mutated (22, 31). Since we did not observe growth phenotypes associated with the *pfp* mutation, we hypothesized that redundant activity or a metabolic bypass might be present. Consequently, we performed insertion-sequencing to identify genes that are non-essential in the wild type but essential in the Δ*pfp* mutant when provided either glucose (glycolytic) or succinate (gluconeogenic) as a carbon source. Notably, we found that disruption of *smc00535* inhibited growth specifically in the Δ*pfp* mutant when grown with succinate (**Appendix S1, Table S2**; **Dataset S2**). The gene *smc00535* is annotated as a putative phosphatase, and thus we hypothesized that this gene encodes a fructose-1,6- bisphosphatase (Fbp). Although fructose-1,6-bisphosphotase activity has been reported for *S. meliloti* (18), a gene for this function has never been reported.

Consistent with our hypothesis, a *smc00535*::pKNOCK mutant displayed wildtype growth on defined media with a variety of carbon sources, whereas a Δ*pfp smc00535*::pKNOCK double mutant could grow with glucose but not with succinate as the carbon source (**Appendix S1, Table S3**). Growth with succinate was restored by complementation with either *pfp* or *smc00535* (**Appendix S1, Table S3**). Moreover, assaying of cell-free extracts demonstrated that the products encoded by *pfp* and *smc00535* both contributed to the overall cellular fructose-1,6-bisphosphotase activity, with loss of both genes yielded this activity undetectable (**Table 1**). Lastly, *in vitro* assays with purified Smc00535 demonstrated a specific activity of 201 nM min^1^ mg^-1^ with F-1,6-biP as a substrate. Together, these results demonstrate that SMc00535 (Fbp) functions as a fructose-1,6- bisphosphatase and is able to compensate for the loss of Pfp in the EMP pathway and allow for continued growth with gluconeogenic substrates.

**Table 1.** Phosphatase activity of *Sinorhizobium meliloti* cell-free extracts.

| Strain | Relevant Genotype | F-1,6-bisphosphatase<br>(nmol min<sup>-1</sup>mg<sup>-1</sup>) * |
| --- | --- | --- |
| Rm1021 | Wild-type | 18 ± 2 |
| SRmD674 | $\Delta pfp::Nm^R$ | 2 ± 0 |
| SRmD727 | *smc00535::pKNOCK* | 6 ± 1 |
| SRmD726 | $\Delta pfp::Nm^R$, *smc00535::pKNOCK* | 0 ± 0 |

\* Specific phosphatase activity in cell-free extracts of *S. meliloti* was measured in the presence of 1 mM fructose-1,6-bisphosphate; values are the averages ± standard deviations of three independent experiments.

### Pfp or Fbp is essential for nitrogen fixation by *S. meliloti* in symbiosis with alfalfa

Mutations in the lower half of the EMP pathway have been shown to affect symbiosis and lead to the formation of ineffective nodules (32). However, strains carrying mutations that resulted in the complete loss of triose phosphate isomerase activity elicited nodules that were effective but had a reduced ability to fix nitrogen (22). To determine the symbiotic phenotypes conveyed by the *pfp* and *fbp* mutations, strains carrying either or both were inoculated onto alfalfa seedlings grown under nitrogen-deficient conditions (**Figure 4**). Mutations in either *pfp* or *fbp* produced no significant reduction in nitrogen fixation compared to the parental strain. On the other hand, mutations in both genes abolished nitrogen fixation, resulting in stunted, chlorotic plants that did not differ significantly in shoot dry weight from the uninoculated control. The nodules produced by these plants were small, white, spherical projections, and lacking zones of differentiation, which is characteristic of ineffective nodules formed by *S. meliloti*. Overall, these results demonstrate that flux through the upper EMP pathway is essential for the symbiotic interaction between *S. meliloti* and alfalfa.

**Figure 4.**
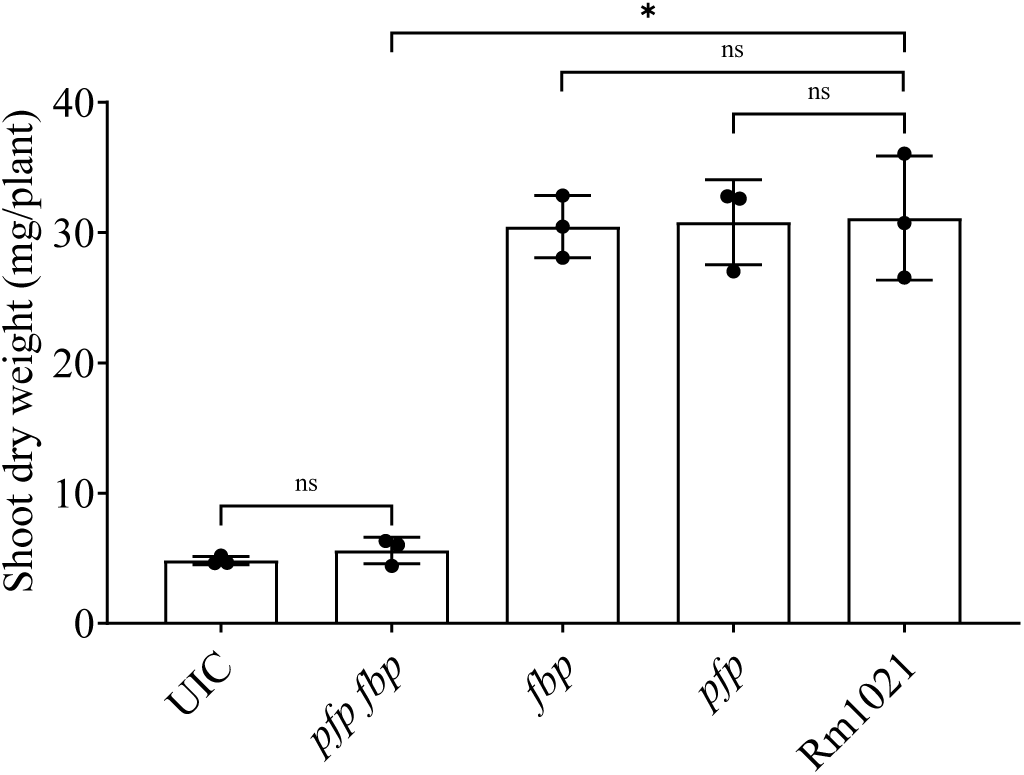
A functional Pfp or Fbp is essential for SNF by *Sinorhizobium meliloti* in symbiosis with alfalfa. Shoot dry weights of alfalfa plants inoculated with various *S. meliloti* strains are shown. Plants were grown in sterile sand-vermiculite media in the absence of nitrogen. Relevant genotypes are indicated on the x-axis. Each bar corresponds to the mean of three biological replicates each containing 10 plants, while the error bars show the standard deviation. Statistically significant differences (*P* < 0.05), as determined by using Student’s t-tests, are indicated with an asterisk (*), while “ns” indicates no statistically significant difference.

### Pfp is widespread amongst phylogenetically diverse rhizobia

Pfk is commonly associated with the EMP pathway and has been referred to as a “textbook enzyme” whereas the presence of Pfp has been primarily associated with plants, archaea, and anaerobic bacteria (24). In contrast, our computational analysis of the phylum *Pseudomonadota* (syn. *Proteobacteria*) revealed that within this phylum, ATP-dependent Pfk is predominately restricted to a monophyletic clade within the class *Gammaproteobacteria* that includes *E. coli*, whereas most other taxa within this phylum that encode a phosphofructokinase instead encode a PPi-dependent Pfp (**Figure 5**; **Appendix S1, Figures S3 and S4**). Notably, most rhizobia of the class *Alphaproteobacteria* (including members of the genera *Sinorhizobium*, *Rhizobium*, *Mesorhizobium*, and *Bradyrhizobium*, among others) were predicted to encode a Pfp homolog, while most members of the family *Rhizobiaceae* (which includes the genera *Sinorhizobium* and *Rhizobium*, among others) were also predicted to encode a Fbp homolog (**Figure 5**; **Appendix S1, Figures S5 and S6**).

**Figure 5.**
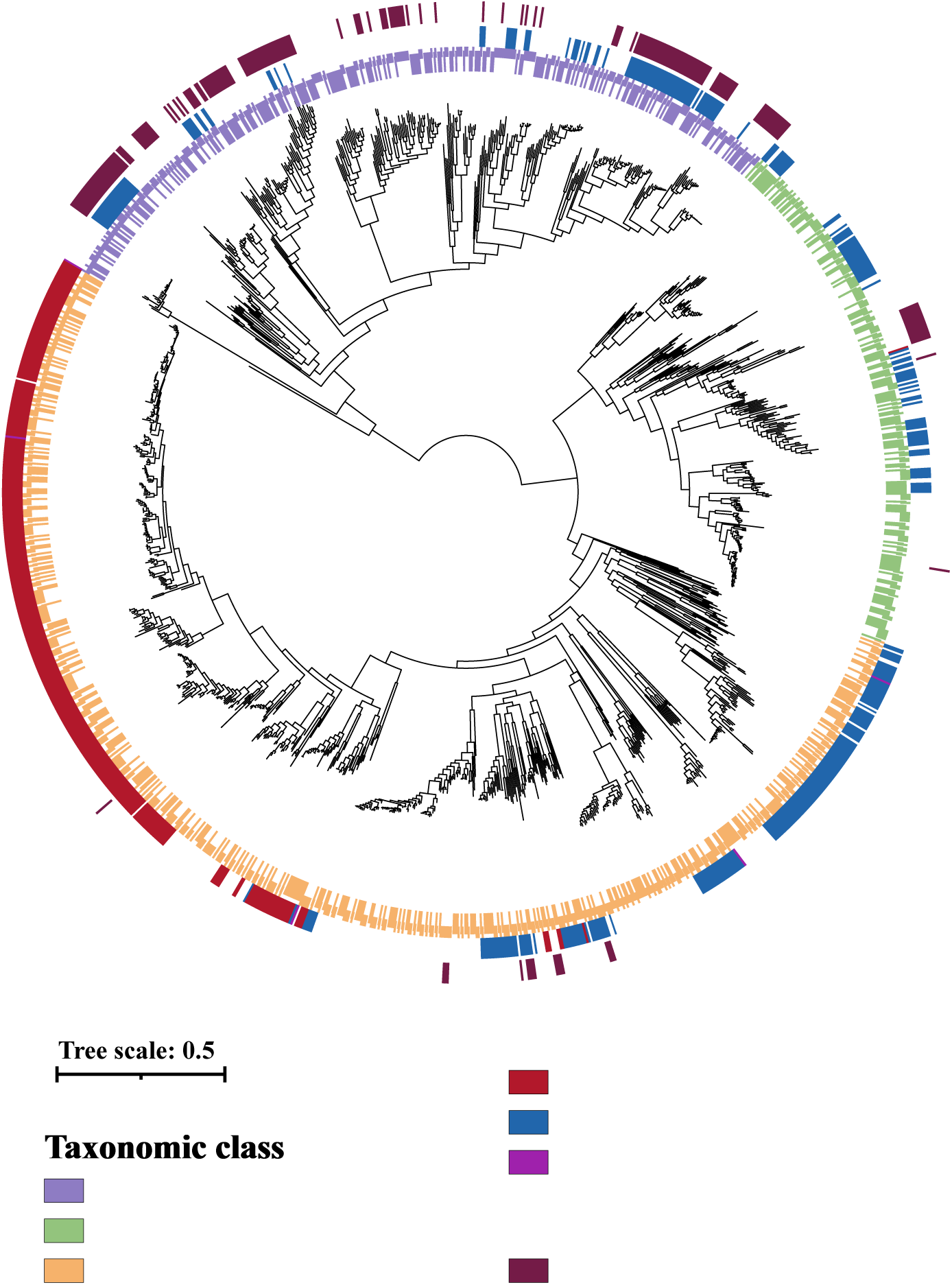
Distribution of phosphofructokinases and fructose-1,6-bisphosphatases in the phylum *Pseudomonadota*. An unrooted, maximum likelihood phylogeny of 1,408 bacteria of the phylum *Pseudomonadota* (syn. *Proteobacteria*) is shown, inferred from the concatenated protein alignments of 27 single copy proteins. The scale bar represents the average number of amino acid substitutions per site. Taxa with long branches were removed for presentation purposes. The three rings around figure represent from inside to outside: (i) taxonomic class (purple – *Alphaproteobacteria*; green – *Betaproteobacteria*; yellow – *Gammaproteobacteria*); (ii) whether the species encodes an ATP-dependent phosphofructokinase (red), a pyrophosphate-dependent phosphofructokinase (blue), both types of enzymes (magenta), or neither (white); and (iii) whether the species encodes (purple) or does not encode (white) an ortholog of the Smc00535 fructose-1,6- bisphosphatase enzyme of *S. meliloti* Rm1021. An interactive version of this phylogeny, with support values, can be accessed at itol.embl.de/shared/Xjwm5Fs4ugTK.

To validate that the computationally predicted Pfp homologs are active, we codon optimized the predicted *pfp* genes from 10 species including both rhizobial and non-rhizobial taxa and introduced them into a *S. meliloti pfp fbp* double mutant. In each case, the introduced gene was able to restore the ability of the mutant to grow on succinate, supporting its annotation as encoding a functional Pfp enzyme (**Appendix S1, Table S3**). Interestingly, introduction of *pfp* genes from *R. etli* or *B. japoncium* resulted in a noticeable impairment in the ability of *S. meliloti* to grow in medium with glucose as the sole carbon source. Overall, our results are consistent with gluconeogenic growth of most alpha-rhizobia involving a PPi-dependent phosphofructokinase.

## DISCUSSION

Phosphofructokinase activity assays of cell-free extracts have previously failed to identify this activity in *S. meliloti*, leading to the erroneous conclusion that the EMP pathway is incomplete in this model legume endosymbiont (10, 33). However, assay conditions have always supplied ATP as a high-energy phosphate donor (27, 28). Here, we show that *S. meliloti* does possesses a complete EMP pathway and that the inability to detect phosphofructokinase activity is because *S. meliloti*, and many other rhizobial species, encodes a PPi-dependent phosphofructokinase rather than an ATP-dependent enzyme. We further show that this activity is redundant due to the presence of a novel frusctose-1,6-bisphosphatase that is widespread in the class *Alphaproteobacteria*; this redundancy would have prevented the identification of these enzymes through classical genetic screens. Lastly, we demonstrate that phosphofructokinase activity or frusctose-1,6-bisphosphatase activity is essential for symbiotic nitrogen fixation with alfalfa.

Pfp was first characterized in the protozoan *Entamobeba histolytica*, where it was shown that the PPi was required as a phosphate donor to phosphorylate F6P, not ATP or GTP (34). Since then, Pfp has been found in bacteria, protists, and plants (35), and the kinetic properties of the *S. meliloti* Pfp are within the ranges of those reported for other bacterial PPi-dependent phosphofructokinases (36–38). However, whereas other Pfp enzymes tend to have a higher affinity for F1,6biP compared to F6P (24), the *S. meliloti* Pfp had similar affinities for both compounds. Nevertheless, the *S. meliloti* Pfp is expected to function only as a F1,6biP diphosphatase *in vivo*; C^13^ labelling studies (12–14), together with gene mutagenesis studies (31, 39), have shown that hexose catabolism in *S. meliloti* is mediated by the ED pathway and that the EMP pathway is unable to complement for the loss of the ED pathway.

In organisms using the EMP pathway for glycolysis, ATP-dependent phosphofructokinases represent a major metabolic checkpoint due to its thermodynamically irreversible nature as well as allosteric inhibition by downstream products such as ADP (40, 41). On the other hand, these mechanisms of protein regulation have not been found in well-characterized Pfp enzymes (24, 40, 42). We therefore speculated that regulation of Pfp may occur at a transcriptional level. In *S. meliloti* and other members of the order *Hyphomicrobiales* (syn. *Rhizobiales*), the LacI-type protein PckR acts as a global regulator of many for many of the genes requires for central carbon metabolism (43, 44). There is no identifiable PckR consensus binding sequence in the upstream promotor region of *pfp*, suggesting it is subject to an alternate mode of transcriptional regulation. However, we note that the promoter region of *pfp* is predicted to contain a Pho box and its expression was shown to be upregulated in a PhoB-dependent fashion under phosphate-limiting growth conditions (45). Considering the regulation of Pfp by PhoB (a central regulator of phosphorus metabolism), and that increased Pfp activity would increase the release of phosphate from phosphorylated intermediates of central carbon metabolism, we hypothesize that *pfp* could be an uncharacterized link between phosphate and carbon metabolism.

It is often thought that within the domain *Bacteria*, *pfp* is predominately limited to anaerobic bacteria (24, 42). It was therefore surprising that in our analysis, PPi-dependent phosphofructokinases were more broadly distributed across the phylum *Pseudomonadota* (syn. *Proteobacteria*) than were ATP-dependent phosphofructokinases, which were instead primarily limited to enteric species of this phylum. It has been debated whether ATP- or PPi-dependent phosphofructokinases are ancestral (24). Given the broader distribution of PPi-dependent enzymes compared to the ATP-dependent enzymes, which were largely found within a monophyletic group of species, our results suggest that the PPi-dependent phosphofructokinases are more ancestral within the phylum *Pseudomonadota*, consistent with recent arguments (24). Our data also supports the notion of frequent changes in donor specificity in this protein family (24), as the PPi- and ATP-dependent phosphofructokinases formed multiple clades in a phylogeny and sequence similarity network, and that PPi-dependent phosphofructokinases are likely more broadly distributed and important than previously recognized.

## MATERIALS AND METHODS

### Bacterial strains, media, and growth conditions

Strains of *S. meliloti* and *E. coli*, as well as plasmids, used for this work are listed in **Appendix S1, Table S4**. *S. meliloti* and *E. coli* were grown at 28°C and 37°C, respectively, with LB or LBmc as complex media along with relevant antibiotic concentrations as previously described (46). Two types of minimal media were used in this work: Vincent’s minimal media (VMM) and MOPS-buffered M9 media (MM9) supplemented with either 15 mM glucose or 15 mM succinate (20 mM for MM9) as the carbon source (31, 47). Genetic manipulations, plasmid constructions, and generation of *S. meliloti* mutant strains were performed as described in **Appendix S1, Text S1**. Oligonucleotides are listed in **Appendix S1, Table S5**.

### Enzyme activity assays

Cell-free extracts and purified proteins were prepared as described in **Appendix S1, Text S2 and S3**. Enzyme activities were measured spectrophotometrically via coupled enzyme reactions. To assay phosphatase activities in mutant strains of *S. meliloti*, crude extracts were added to a reaction buffer consisting of 100 mM MOPS pH 8, 5mM MgCl_2_, 15 mM NaH_2_PO_4_ 2mM PPi, 3.5 U/mL phosphoglucose isomerase (Roach), 3.5 U/mL glucose-6-phosphate dehydrogenase (Roach), 0.3 mM NADP^+^ (Sigma-Millipore). Reactions were initiated through the addition of 5 mM frucose- 1,6-bisphosphate. Specific activities are reported as a mean of three technical triplicates.

The kinetics of Pfp were characterized for both the forward and reverse reactions by coupling its activity to the oxidation of NADH and reduction of NADP+, respectively. The forward reaction buffer contained 100 mM MOPS pH 8, 5mM MgCl_2_, 2 mM PPi, 10U/mL rabbit aldolase (Sigma), triosphosphate isomerase (Sigma), 5U/mL glycerol-3-phosphate dehydrogenase (Roach), 2 mM NADH, and purified Pfp. The forward reaction was initiated by adding fructose-6- phosphate. The reverse reaction buffer consisted of 100 mM MOPS pH 8, 5mM MgCl_2_, 15 mM NaH_2_PO_4_, 2mM PPi, 3.5 U/mL phosphoglucose isomerase, 3.5 U/mL glucose-6-phosphate dehydrogenase, 3 mM NADP^+^ and purified Pfp. The reverse reaction was initiated through the addition of fructose-1,6-phosphate. The affinity of Pfp for PPi was measured similarly to the forward reaction, except PPi was added at varying concentrations to initiate a reaction. Specific activities are representative of an average of at least three technical triplicates measured at 340 nm. Michaelis-Menton kinetics were calculated by linear regression analysis using GraphPad Prism 9.3.1.

The specific activity of Fbp was measured by coupling the release of orthophosphate to uric acid production as described by Guillen Suarez *et al.* (48). Reactions were initiated with the addition of 5 mM F1,6biP and activity recorded at 293 nm. Specific activity of Fbp is reported in nmol of uric acid produced per minute per mg of purified Fbp. The assay was performed in three technical triplicates.

### Metabolite analysis

Cultures of *S. meliloti* were grown in 500 mL VMM-glucose or VMM-succinate overnight with shaking. After reaching an OD_600_ ∼ 0.5, 100 mL volumes of each culture were harvested by centrifugation, washed with double distilled water, and flash frozen in liquid N_2_ before weighing. Metabolite extraction and LC-MS was performed at the Metabolomics Innovation Resource (MIR) -McGill University as described elsewhere (49). All data represents an average of 3 biological replicates.

Metabolite concentrations were normalized by log-transformation. A significant difference was determined by performing Welch’s two sample t-test with a P-value threshold of 0.05. The heatmap was generated using the z-scores of each metabolite concentration relative to the mean of log-transformed concentrations for each metabolite across all strains and growth conditions. Principle component analysis was performed with the pheatmap package in R-studio.

### Symbiosis assays

Symbiosis assays were performed as described previously (49). Briefly, surface sterilized alfalfa seedings were germinated on water agar and transplanted to a sterile 1:1 mixture of sand and vermiculite soaked with N-free Jensen’s media in Leonard jar assemblies (∼10 plants/pot) (50). Plants were inoculated with ∼10^7^ CFU/mL and grown for 28-33 days at 25 °C with a 16 h day light cycle. Plant tissue above the cotyledons were harvested and dried to calculate shoot dry weights. Results are reported as the mean of at least three independent biological replicates.

### Insertion-sequencing (IN-seq)

IN-seq libraries were prepared for both wildtype *S. meliloti* and the Δ*pfp* mutant using pSAM_Rl (Perry and Yost, 2014) as described in **Appendix S1, Text S4**. IN-seq sequencing libraries were prepared as described in **Appendix S1, Text S5**, and sequenced at The Centre for Applied Genomics (Toronto, Canada) using one lane of a SP flow cell on an Illumina NovaSeq 6000 instrument and 25% PhiX spike-in, to generate 100 bp single-end reads. Data analysis was performed as described in **Appendix S1, Text S6**.

### Phylogenetic distribution of Pfp and Fbp orthologs

The RefSeq proteomes of 1,475 representative proteobacteria (**Dataset S3**) were downloaded from the National Center for Biotechnology Information (NCBI) database on 30 June 2022. A phylogeny of the 1,475 species was constructed using a multilocus sequence analysis (MLSA) approach following our previously defined pipeline (diCenzo *et al*, 2019) and as described in **Appendix S1, Text S7**. Phosphofructokinases (**Dataset S4**) and fructose-1,6-bisphosphatases (**Dataset S5**) were identified in the representative proteobacteria using hidden Markov models (HMMs) with a modified reciprocal hit approach as described in **Appendix S1, Text S8 and S9**. Substrate preferences of the phosphofructokinases were determined using HMMs as described in **Appendix S1, Text S8**.

## Data availability

Raw Illumina data are available from the NCBI Short Read Archive (SRA) database under BioProject accession PRJNA1468229. Scripts to repeat all computational analyses are available at github.com/diCenzo-Lab/020_2026_Sinorhizobium_meliloti_pfp. Phylogenies are available through iTOL at itol.embl.de/shared/Xjwm5Fs4ugTK.

## ACKNOWLEDGEMENTS

We thank Derek Kim for his contributions in making SRmD674, Dr. Justin Hawkins for assisting with the enzyme assays, and Kiera Antaya for assistance with the insertion-sequencing. This work was funded through the Natural Sciences and Engineering Research Council of Canada (NSERC) Discovery Grants program through grants to IJO (RGPIN-2024-05461) and GCD (RGPIN-2020- 0700). This research was enabled in part by support provided by Compute Ontario (computeontario.ca) and the Digital Research Alliance of Canada (alliancecan.ca).

## REFERENCES

1. D. S. LeBauer, K. K. Treseder, Nitrogen limitation of net primary productivity in terrestrial ecosystems is globally distributed. Ecology 89, 371–379 (2008).

2. V. Smil, Feeding the world: a challenge for the twenty-first century (MIT Press, 2000).

3. S. Ghavam, M. Vahdati, I. A. G. Wilson, P. Styring, Sustainable ammonia production processes. *Front*. Energy Res. 9 (2021).

4. T. E. Crews, M. B. Peoples, Legume versus fertilizer sources of nitrogen: ecological tradeoffs and human needs. Agric. Ecosyst. & Environ. 102, 279–297 (2004).

5. S. A. Montzka, E. J. Dlugokencky, J. H. Butler, Non-CO2 greenhouse gases and climate change. Nature 476, 43–50 (2011).

6. Y. Gao, A. Cabrera Serrenho, Greenhouse gas emissions from nitrogen fertilizers could be reduced by up to one-fifth of current levels by 2050 with combined interventions. Nat. Food. 4, 170–178 (2023).

7. D. A. Day, P. S. Poole, S. D. Tyerman, L. Rosendahl, Ammonia and amino acid transport across symbiotic membranes in nitrogen-fixing legume nodules. Cell. Mol. Life Sci. 58, 61–71 (2001).

8. A. Wen, et al., Enabling biological nitrogen fixation for cereal crops in fertilized fields. ACS Synth. Biol. 10, 3264–3277 (2021).

9. K. M. Jones, H. Kobayashi, B. W. Davies, M. E. Taga, G. C. Walker, How rhizobial symbionts invade plants: the *Sinorhizobium*–*Medicago* model. Nat. Rev. Microbiol. 5, 619– 633 (2007).

10. B. A. Geddes, I. J. Oresnik, Physiology, genetics, and biochemistry of carbon metabolism in the alphaproteobacterium *Sinorhizobium meliloti*. Can. J. Microbiol. 60, 491–507 (2014).

11. G. C. Dicenzo, et al., Multidisciplinary approaches for studying rhizobium–legume symbioses. Can. J. Microbiol. 65, 1–33 (2019).

12. T. Fuhrer, E. Fischer, U. Sauer, Experimental identification and quantification of glucose metabolism in seven bacterial species. J. Bacteriol. 187, 1581–1590 (2005).

13. J. C. Portais, P. Tavernier, I. Gosselin, J. N. Barbotin, Cyclic organization of the carbohydrate metabolism in *Sinorhizobium meliloti*. Eur. J. Biochem. 265, 473–480 (1999).

14. I. Gosselin, O. Wattraint, D. Riboul, J.-N. Barbotin, J.-C. Portais, A deeper investigation on carbohydrate cycling in *Sinorhizobium meliloti*. FEBS Lett. 499, 45–49 (2001).

15. N. J. Booth, P. M. C. Smith, S. A. Ramesh, D. A. Day, Malate Transport and metabolism in nitrogen-fixing legume nodules. Molecules 26, 6876 (2021).

16. T. M. Finan, J. M. Wood, D. C. Jordan, Symbiotic properties of C4-dicarboxylic acid transport mutants of *Rhizobium leguminosarum*. J.Bacteriol. 154, 1403–1413 (1983).

17. J. Prell, P. Poole, Metabolic changes of rhizobia in legume nodules. Trends Microbiol. 14, 161–168 (2006).

18. T. M. Finan, I. Oresnik, A. Bottacin, Mutants of *Rhizobium meliloti* defective in succinate metabolism. J. Bacteriol. 170, 3396–3403 (1988).

19. B. T. Driscoll, T. M. Finan, NAD+ -dependent malic enzyme of *Rhizobium meliloti* is required for symbiotic nitrogen fixation. Mol. Microbiol. 7, 865–873 (1993).

20. B. T. Driscoll, T. M. Finan, NADP+-dependent malic enzyme of *Rhizobium meliloti*. J. Bacteriol. 178, 2224–2231 (1996).

21. M. F. Dunn, Tricarboxylic acid cycle and anaplerotic enzymes in rhizobia. FEMS Microbiol. Rev. 22, 105–123 (1998).

22. N. J. Poysti, E. D. M. Loewen, Z. Wang, I. J. Oresnik, *Sinorhizobium meliloti* pSymB carries genes necessary for arabinose transport and catabolism. Microbiology 153, 727–736 (2007).

23. S. Kaur, J. P. Hawkins, I. J. Oresnik, Suppression of a transketolase mutation leads to only partial restoration of symbiosis in *Sinorhizobium meliloti*. Mol. Plant Microbe Interact. 38, 505–517 (2025).

24. J. A. Compton, W. M. Patrick, The more we learn, the more diverse it gets: structures, functions and evolution in the Phosphofructokinase Superfamily. Biochem. J. 482, 467–483 (2025).

25. T. Schirmer, P. R. Evans, Structural basis of the allosteric behaviour of phosphofructokinase. Nature 343, 140–145 (1990).

26. D. Capela, et al., Analysis of the chromosome sequence of the legume symbiont *Sinorhizobium meliloti* strain 1021. Proc. Natl. Acad. Sci. U.S.A. 98, 9877–9882 (2001).

27. J. J. Irigoyen, M. Sanchez-Diaz, D. W. Emerich, Carbon metabolism enzymes of *Rhizobium meliloti* cultures and bacteroids and their distribution within alfalfa nodules. Appl. Environ. Microbiol. 56, 2587–2589 (1990).

28. A. Arias, C. Cervenansky, A. Gardiol, G. Martinez-Drets, Phosphoglucose isomerase mutant of *Rhizobium meliloti*. J. Bacteriol. 137, 409–414 (1979).

29. G. C. diCenzo, M. Tesi, T. Pfau, A. Mengoni, M. Fondi, Genome-scale metabolic reconstruction of the symbiosis between a leguminous plant and a nitrogen-fixing bacterium. Nat. Commun. 11, 2574 (2020).

30. M. N. Price, et al., Mutant phenotypes for thousands of bacterial genes of unknown function. Nature 557, 503–509 (2018).

31. G. C. diCenzo, T. M. Finan, Genetic redundancy is prevalent within the 6.7 Mb *Sinorhizobium meliloti* genome. Mol. Genet. Genomics. 290, 1345–1356 (2015).

32. T. M. Finan, E. McWhinnie, B. Driscoll, R. J. Watson, Complex symbiotic phenotypes result from gluconeogenic mutations in *Rhizobium meliloti*. Mol. Plant Microbe Interact. (1991).

33. M. D. Stowers, Carbon metabolism in *Rhizobium* species. Annu. Rev. Microbiol. 39, 89–108 (1985).

34. R. E. Reeves, D. J. South, H. J. Blytt, L. G. Warren, Pyrophosphated-fructose 6-phosphate 1- phosphotransferase. J. Biol. Chem. 249, 7737–7741 (1974).

35. E. Mertens, Pyrophosphate-dependent phosphofructokinase, an anaerobic glycolytic enzyme? FEBS Lett. 285, 1–5 (1991).

36. C. Pfleiderer, J.-H. Klemme, Pyrophosphat-abhängige ᴅ-Fructose-6-phosphat Phosphotransferase in *Rhodospirillaceae* / Pyrophosphate-dependent ᴅ-Fructose-6- phosphate-phosphotransferase in *Rhodospirillaceae*. Z. Naturforsch. C 35, 229–238 (1980).

37. M. Frese, S. Schatschneider, J. Voss, F.-J. Vorhölter, K. Niehaus, Characterization of the pyrophosphate-dependent 6-phosphofructokinase from *Xanthomonas campestris* pv. campestris. Arch. Biochem. Biophys. 546, 53–63 (2014).

38. O. N. Rozova, V. N. Khmelenina, S. Vuilleumier, Y. A. Trotsenko, Characterization of recombinant pyrophosphate-dependent 6-phosphofructokinase from halotolerant methanotroph *Methylomicrobium alcaliphilum* 20Z. Res. Microbiol. 161, 861–868 (2010).

39. G. C. diCenzo, et al., Robustness encoded across essential and accessory replicons of the ecologically versatile bacterium *Sinorhizobium meliloti*. PLoS Genet. 14, undefined- undefined (2018).

40. A. M. C. R. Alves, G. J. W. Euverink, H. Santos, L. Dijkhuizen, Different physiological roles of ATP- and PPi-dependent phosphofructokinase isoenzymes in the methylotrophic *Actinomycete Amycolatopsis methanolica*. J. Bacteriol.183, 7231–7240 (2001).

41. E. Hofmann, The significance of phosphofructokinase to the regulation of carbohydrate metabolism. Rev. Physiol. Biochem. Pharmacol. 75, 1–68 (1976).

42. B. Siebers, H.-P. Klenk, R. Hensel, PPi-Dependent phosphofructokinase from *Thermoproteus tenax*, an archaeal descendant of an ancient line in phosphofructokinase evolution. J. Bacteriol. 180, 2137–2143 (1998).

43. M. Østerås, S. A. P. O’Brien, T. M. Finan, Genetic analysis of mutations affecting *pckA* regulation in *Rhizobium* (*Sinorhizobium*) *meliloti*. Genetics 147, 1521–1531 (1997).

44. G. C. diCenzo, Z. Muhammed, M. Østerås, S. A. P. O’Brien, T. M. Finan, A key regulator of the glycolytic and gluconeogenic central metabolic pathways in *Sinorhizobium meliloti*. Genetics 207, 961–974 (2017).

45. Z.-C. Yuan, R. Zaheer, R. Morton, T. M. Finan, Genome prediction of PhoB regulated promoters in *Sinorhizobium meliloti* and twelve proteobacteria. Nucleic Acids Res. 34, 2686– 2697 (2006).

46. J. Sambrook, E. F. Fritsch, T. Maniatis, Molecular cloning: A laboratory manual (Cold Spring Harbor Laboratory, 1989).

47. J. M. Vincent, A manual for the practical study of root-nodule Bacteria ([Published for the] International Biological Programme [by] Blackwell Scientific, 1970).

48. A. S. G. Suárez, A. Stefan, S. Lemma, E. Conte, A. Hochkoeppler, Continuous enzyme- coupled assay of phosphate- or pyrophosphate-releasing enzymes. Biotechniques 53, 99–103 (2012).

49. J. P. Hawkins, P. A. Ordonez, I. J. Oresnik, Characterization of mutations That affect the nonoxidative pentose phosphate pathway in *Sinorhizobium meliloti*. J. Bacteriol. 200, e00436–17 (2018).

