## Appendix S1 for "*Sinorhizobium meliloti* possesses a complete Embden-Meyerhoff-Parnas pathway that is indispensable for symbiotic nitrogen fixation"

**This PDF file includes:**

Text S1 to S9 (pages 2-6)
Tables S1 to S5 (pages 7-12)
Figures S1 to S7 (pages 13-19)
Legends for Datasets S1 to S5 (pages 20-21)
SI References (pages 22-24)

**Text S1. Genetic manipulations, plasmid constructions, and mutations**

Plasmid isolation, ligations, restrictions, transformations, and agarose gel electrophoresis were performed as previously described (1). Conjugations and generalized transductions were carried out with the *E. coli* MT616 helper strain and ΦM12 bacteriophage, respectively (2, 3). Complementing plasmids were constructed using the ORFeome library (4, 5). Knockout mutations of *pfp* or *fbp* were made using the pKNOCK series of vectors carrying homologous DNA sequences flanking the target gene amplified by PCR with high-fidelity Q5 DNA polymerase (NEB) (6). An allelic exchange of *pfp* with a neomycin resistance gene was generated via double crossover recombination with the suicide vector pJQ200SK+ carrying the Nm cassette with adjacent flanks homologous to the target locus (7). An unmarked deletion of *pfp* was made using the same strategy without a Nm cassette. All mutations were confirmed by DNA sequencing. Functional complementation of *pfp and pfkA* orthologs in *S. meliloti* was performed with codon optimized phosphfructokinase genes (purchased from Biobasic Inc.) of representative proteobacterial species identified by phylogenetic analysis.

**Text S2. Preparation of cell-free extracts**

Rhizobial strains used in crude enzyme activity assays were grown overnight in 2 L volumes. Once an optical density at 600 nm ($OD_{600}$) of > 1.0 was reached, rhizobia were harvested via centrifugation followed by resuspension in lysis buffer containing 100 mM Tris, 2 mM $MgCl_2$, 1 mM dithiothreitol (DTT), and 100 mM PMSF. Cells were lysed via passage through an AVESTIN emulsiflex at ~20,000 psi. Cell-free extracts were isolated and stored at -80˚C. The concentration of cell-free lysates were measured as described by Bradford (8).

**Text S3. Protein expression and purification**

The *pfp* ORF in *S. meliloti* Rm1021 was codon optimized for overexpression in an *E. coli* JM109 background by Genscript (Piscataway, New Jersey, USA). The optimized gene was amplified by PCR and cloned into the pQE-9 overexpression vector in frame with an N-terminal RGS-His6X polyhistidine tag. The *fbp* gene was codon optimized in a similar manner and was synthesized in the expression vector pQE-30 in frame with an N-terminal RGS-His6X polyhistidine tag by Bio Basic Inc (Markham, Ontario). Both expression constructs were transformed into *E. coli* JM109. Overexpression and purification were protocols were adapted from Kohlmeier *et al*. 2021. Cultures were induced with 1 mM of IPTG at an $OD_{600}$ ~ 0.4 and grown two hours followed by lysis with an AVESTIN emulsiflex at ~20,000 psi. Cell-free lysates were stored at –80˚C. Proteins were purified via immobilized metal affinity chromatography (IMAC) using $Ni^{2+}$-bound Sepharose His SpinTrap (cytiva) and dialyzed overnight in 50 mM Tris pH 8, 0.1 mM EDTA, 1 mM DTT at 4˚C. Purification fractions were visualized by SDS-PAGE and Western blot analysis (9, 10) (Figures S1 and S7). The concentration of purified proteins was quantified using the Bradford method (8).

**Text S4. Insertion-sequencing (Tn-seq) experimental setup**

Seven replicate cultures of each of wildtype *S. meliloti* and the Δ*pfp* mutant (SRmD673) were grown overnight in LBmc. In parallel, nine replicate cultures of *E. coli* SM10λpir (pSAM_Rl) (11) were grown in LB Amp; these cultures were diluted 1:100 the next morning into fresh LB Amp and incubated until late afternoon. Next, 14 replicate mating spots were prepared for each *S. meliloti* strain and incubated at 30˚C for ~24 hours, after which each mating spot was resuspended and washed in 0.85% saline. Half of each mating spot was spread plated on MM9-succinate with streptomycin (Sm) and neomycin (Nm) while the other half was plated on MM9-glucose Sm Nm and incubated at 30˚C for ~16 hours. All bacteria from each plate were scrapped off the surface and cell suspensions from the same treatment were combined. For each *S. meliloti* strain, 50 mL cultures of MM9-glucose Sm Nm and MM9-succinate Sm Nm were inoculated to a final $OD_{600}$ of ~0.05 and incubated at 30˚C for ~24 hours with shaking (MM9-succinate) or ~27.5 hours (MM9-glucose). Cultures were pelleted and resuspended with saline. The equivalent of 10 mL at an $OD_{600}$ of 1 were transferred to a new tube, centrifuged at 16,000 x *g*, and pelleted stored at -20˚C until use.

**Text S5. IN-seq DNA library preparation and sequencing**

IN-seq DNA library preparation was performed using an adaptation of a previously described protocol (12). Total genomic DNA was isolated from each of the IN-seq cell pellets using phenol – chloroform extractions followed by ammonium acetate – isopropanol precipitations as described elsewhere (13). RNase A treated DNA pellets were then resuspended in nuclease-free water. Approximately 600 ng of gDNA per sample was fragmented using the NEBNext dsDNA fragmentase (NEB) for 17 minutes at 37˚C according to the manufacturer's protocol. Fragmented samples were purified using Monarch PCR & DNA Cleanup kits (NEB) and eluted in nuclease-free water. Short oligo-C tails were added to the fragmented gDNA using terminal transferase (NEB) for 30 minutes at 37˚C according to the manufacturer's protocol with the following modifications: the entire DNA suspensions, 0.4 µL of enzyme, and 1.5 µL of 1 mM dCTP were added to each 50 µL reaction. Tailed samples were purified using Monarch PCR & DNA Cleanup kits (NEB), eluted in EB buffer, and stored at -20˚C until use.

DNA samples were diluted to 10 ng/µL and PCRs were performed using Q5 polymerase and the primers 1TN-mariner and 1GG (Table S5), according to the manufacturer's protocol. Triplicate reactions were performed per sample, with 50 ng of DNA added to each PCR in 25 µL reaction volumes. Twenty-five PCR cycles were performed with each cycle consisting of 98˚C for 10 s, 61˚C for 15 s, and 72˚C for 30 s. Samples corresponding to the same DNA sample were pooled, purified using Monarch PCR & DNA Cleanup kits (NEB), and eluted in EB buffer. The PCR products were subsequently used as templates in a second round of PCRs. For each sample, triplicate 25 µL reactions were set-up, each using 50 ng of template, the PCR cycling conditions described above, an equimolar mix of 2TNA-mariner, 2TNB-mariner, and 2TNC-mariner as the forward primer, and one of 2BAR5, 2BAR6, 2BAR7, or 2BAR8 as the reverse primer (Table S5). Samples corresponding to the same DNA sample were pooled, purified using Monarch PCR & DNA Cleanup kits (NEB), eluted in EB buffer, and stored at -20˚C until use.

Approximately 1.2 µg of each PCR product were combined and mixed with $1/3^{rd}$ volume of 3 M sodium acetate and 1.5 volume of isopropanol, and then incubated at -20˚C for 30 minutes. DNA was pelleted by centrifugation, washed once with 70% ethanol and once with 90% ethanol, dried, and then resuspended in 32 µL of T10E1 buffer (10 mM Tris-HCl, 1 mM EDTA, pH 8.0). The full sample was then run on a BluePippin instrument with a 2% agarose gel, and fragments between 400 and 600 bp were recovered. The combined sample was then sequenced at The Centre for Applied Genomics (Toronto, Canada) using one lane of a SP flow cell on an Illumina NovaSeq 6000 instrument and 25% PhiX spike-in, to generate 100 bp single-end reads.

**Text S6. IN-seq data analysis**

BBduk version 38.96 (14) was used to remove contamination from the raw Illumina reads. Following this, the reads were trimmed using Trimmomatic version 0.39 (15). A custom Perl script was used to identify trimmed reads containing the sequence corresponding to the end of the mariner transposon and extract just the sequence downstream of the transposon end, after which BBduk was used to discard reads shorter than 18 nt. The extracted reads were next aligned to the *S. meliloti* Rm1021 genome sequence (NCBI RefSeq Accession GCF_000006965.1) (16) using bowtie2 version 2.4.5 (17) and the output sorted using samtools version 1.5.1-33-g906f657 (18). Custom Perl and Matlab (R2019a) scripts were then used to identify all TA sites in the *S. meliloti* Rm1021 genome, link individual TA sites to genes, and count the number of TA sites per gene.

Custom Matlab (R2019a) scripts were used to calculate two summary metrics for comparison of gene fitness results across conditions. (i) A normalized read count was calculated for each gene by dividing the number of reads mapping to a specific gene by the total number of mapped reads divided by one million. (ii) A normalized read count per TA site was calculated by dividing the number of reads mapping to a specific TA site by the total number of mapped reads and then multiplying by one million. Next, for each gene, the median normalized read count per TA site within a given gene was calculated, excluding TA sites with no reads. Lastly, the median normalized read count per TA site for a given gene was multiplied by the number of TA sites in the gene with at least one mapped read, and then divided by the total number of TA sites in the gene (regardless of whether reads mapped to it).

**Text S7. Species phylogenetic analyses**

The RefSeq proteomes of 1,475 representative proteobacteria (Dataset S2) were downloaded from the National Center for Biotechnology Information (NCBI) database on 30 June 2022. A phylogeny of the 1,475 species was constructed using a multilocus sequence analysis (MLSA) approach following our previously described pipeline (19). Briefly, the AMPHORA2 pipeline (20) was paired with custom scripts to extract a set of 27 single-copy proteins (Frr, InfC, NusA, RplA, RplB, RplC, RplD, RplE, RplF, RplK, RplL, RplM, RplN, RplP, RplS, RplT, RpmA, RpoB, RpsB, RpsE, RpsI, RpsJ, RpsK, RpsM, RpsS, SmpB, Tsf) found in at least 98% of the proteomes. Orthologous groups were aligned using MAFFT 7.453(21), following which the alignments were trimmed using TRIMAL 1.4rev22 with the automated1 option (22). Alignments were then concatenated for use in constructing a maximum-likelihood (ML) phylogeny. First, the concatenated protein alignment was used as input for ModelFinder (23) as implemented in IQ-TREE version 2.2.2.4 (24), and the best scoring model was identified based on Bayesian information criterion (BIC). IQ-TREE was then used to infer ML phylogenies from the concatenated alignment using the best-scoring model (LG+I+R10). Branch supports were assessed in IQ-TREE using the Shimodaira-Hasegawa-like approximate likelihood ratio test (SH-aLRT) (25) and ultrafast jackknife analysis with a subsampling proportion of 40%, with both metrics calculated from 1000 replicates. All phylogenies created in this study were visualized with the iTOL web server (26).

**Text S8. Identification and phylogenetic analysis of phosphofructokinase proteins**

All hidden Markov models (HMMs) from the Pfam version 36.0 (27) and TIGRFAM version 15.0 (28) databases were downloaded on 20 January 2024. In addition, alignments for all NCBI Conserved Domains (CD) and Cluster of Orthologous Genes (COG) profiles were downloaded (20 January 2024) and used to generate HMMs with the hmmbuild function of HMMER version 3.3 (29). All HMMs were concatenated into a single database and prepared for use with the hmmpress function of HMMER.

The PF00365 seed alignment of Pfam and the TIGR02045, TIGR02477, TIGR02482, and TIGR02483 seed alignments of TIGRFAM were downloaded and used to generate HMMs with the hmmbuild function of HMMER. Using the hmmsearch function of HMMER, each HMM was searched against a concatenated proteome of the 1,475 representative proteobacteria, and all hits were extracted. The extracted proteins were screened against the concatenated HMM database using the hmmscan function of HMMER, and the top-scoring HMM of each protein was recorded. The 846 proteins whose top hit were cd00763, COG0205, or any of the five HMMs listed above, were collected as putative phosphofructokinases; five proteins were removed due to them having >20% gaps in a preliminary alignment of the proteins, leaving a set of 841 putative phosphofructokinases. To predict the substrate preference of these enzymes, the hmmscan function of HMMER was used to screen the 841 proteins against the TIGR02482 (ATP), TIGR02477 (PPi), TIGR02483 (PPi), and TIGR02045 (ADP) HMMs, and substrate preference assigned according to the top scoring HMM.

The 841 predicted phosphofructokinase enzymes were aligned using MAFFT and the alignment trimmed using TRIMAL. IQ-TREE was then used to infer a ML phylogeny from the trimmed alignment using the LG+I+R10 model. The LG+I+R10 model was used as it was identified as the best-scoring model by the IQ-TREE implementation of ModelFinder, with model search limited to the LG, WAG, JTT, Q.pfam, JTTDCMut, DCMut, VT, PMB, BLOSUM62, and Dayhoff models. Branch supports were assessed in IQ-TREE using SH-aLRT and an ultrafast bootstrap analysis, with both metrics calculated from 1000 replicates.

**Text S9. Identification and phylogenetic analysis of fructose-1,6-bisphosphatases**
The seed alignments of PF00316 and PF03320 of Pfam, TIGR00330 of TIGRFAM, COG0158 and COG0483 of COG, and cd01515 of NCBI CD were downloaded and used to generate HMMs with the hmmbuild function of HMMER. As described for phosphofructokinases (Text S8), each HMM was searched against the concatenated proteobacterial proteome, and top hits searched against the concatenated HMM database. The 2,864 proteins whose top hit were cd00354, cd01516, cd01517, or any of the six HMMs listed above were collected as putative fructose-1,6-bisphosphatases. The 2,864 proteins were filtered to remove proteins whose length was ≤50 amino acids or ≥400 amino acids, leaving 2,848 proteins. One additional protein was removed as its inclusion resulted in a short alignment following trimming during a preliminary alignment of the proteins, leaving a final set of 2,847 putative fructose-1,6-bisphosphatases. The 2,847 predicted fructose-1,6-bisphosphatases were aligned using MAFFT and the alignment trimmed using TRIMAL. IQ-TREE was then used to infer a ML phylogeny from the trimmed alignment using the Q.pfam+R10 model, as this model was identified as the best-scoring model by ModelFinder. Branch supports were assessed in IQ-TREE using SH-aLRT and an ultrafast bootstrap analysis, with both metrics calculated from 1000 replicates.

Two approaches were taken to further classify fructose-1,6-bisphosphatases as putative Smc00535 orthologs. First, the 232 proteins with cd01517 and COG0483 as the top two scoring HMMs in the hmmscan search were classified as putative Smc00535 orthologs. Together with an additional 19 proteins, these 232 proteins formed a well-supported monophyletic group in the fructose-1,6-bisphosphatases ML phylogeny; as a result, we included these additional 19 proteins as putative Smc00535 orthologs, giving a final set of 251 proteins. Second, a sequence similarity network (SSN) was constructed for the 2,847 putative fructose-1,6-bisphosphatases using the online Enzyme Function Initiative – Enzyme Similarity Tool (EFI-EST; efi.igb.illinois.edu/efi-est/) (30, 31) using default settings and an alignment score threshold of 48. The resulting SSN was visualized using Cytoscape version 3.10.1 (32), and a cluster of 251 proteins that included Smc00535 was identified. As the 251-protein Smc00535 cluster in the SSN was identical to the

251-protein clade identified in the fructose-1,6-bisphosphatases ML phylogeny, we annotated these 251 proteins as putative SMc00535 orthologs.

**Table S1.** Normalized concentration of intracellular metabolites.

| **Metabolite (nM/mg)** | **Rm1021 glucose** | **Rm1021 succinate** | **Δ*pfp* glucose** | **Δ*pfp* succinate** |
|---|---|---|---|---|
| **ED/EMP** | | | | |
| G6P/G6P/G1P | 1943 | 2772 | 2027 | 5153 |
| Gluconate | 212 | 68 | 154 | 19** |
| 6P-Gluconic acid | 321 | 304 | 718* | 643 |
| 2P-Glyceric acid/3P-Glyceric acid | 95 | 386 | 398 | 1434** |
| Phosphoenolpyruvate | 298 | 258 | 290 | 374 ** |
| F1,6 biP | 126 | 95 | 2011* | 979** |
| **PPP** | | | | |
| Ribose 5P/Ribulose 5P | 731 | 581 | 882 | 918 |
| Sedoheptulose-7P | 441 | 1971 | 562 | 1871 |
| 2-Deoxy-Ribose-5P | 82 | 74 | 189 | 426 |
| Ribose | 19682 | 15279 | 27800 | 13394** |
| **TCA** | | | | |
| Isocitrate | 154 | 86 | 86* | 50 |
| Fumarate | 732 | 703 | 526 | 1015 |
| Cis-aconitic acid | 14 | 12 | 15 | 12 |
| α-ketoglutrate | 73 | 29 | 41 | 30 |
| 2-Hydroxyglutarate | 27 | 15 | 14* | 12 |
| Succinate | 1295 | 47393 | 3831 | 29963** |
| Lactate | 458 | 369 | 386 | 228 |
| Malate | 304540 | 234833 | 195822 | 206870 |

Values represent the mean of triplicate samples.

* Significant differences in metabolite concentrations of glucose grown cells when compared to the glucose grown wild type.

** Significant differences in metabolite concentrations of succinate grown cells when compared to the succinate grown wild type. Significance ($P < 0.05$) was determined by Student's T-test.

**Table S2.** Insertion-seq data for genes of interest.

| **Locus tag** | Gene name | Rm1021 glucose | Rm1021 succinate | Δ*pfp* glucose | Δ*pfp* succinate |
|---|---|---|---|---|---|
| *smc03070* | *zwf* | 1 | 103 | 2 | 250 |
| *smc02562* | *pckA* | 287 | 4 | 74 | 3 |
| *smc01852* | *pfp* | 309 | 11 | NA* | NA* |
| *smc00535* | *fbp* | 385 | 291 | 113 | 1 |

Numbers represent normalized read counts per gene, with lower numbers representing the gene was more critical for growth in the given strain/environment.
* Gene was replaced with $Nm^R$ cassette in the Δ*pfp* strain.

**Table S3.** Functional complementation of a *Sinorhizobium meliloti* Δ*pfp fbp*::pKnock mutant with codon optimized *pfp* orthologs from diverse proteobacteria.

| **Strain** | **Relevant Genotype** | **Source of *pfp*** | **Growth with Carbon Source *** | | |
|---|---|---|---|---|---|
| | | | **Glc** | **Suc** | **LB** |
| **Rm1021** | Wild type | NA† | ++ | ++ | ++ |
| **SRmD674** | Δ*pfp*::$Nm^R$ | NA† | ++ | ++ | ++ |
| **SRmD727** | *fbp*::pKnock $Gm^R$ | NA† | ++ | ++ | ++ |
| **SRmD726** | Δ*pfp*::$Nm^R$, *fbp*::pKnock | NA† | ++ | - | ++ |
| **SRmD726** | Δ*pfp*::$Nm^R$ *fbp*::pKnock | *S. meliloti* | ++ | ++ | ++ |
| **SRmD726** | Δ*pfp*::$Nm^R$ *fbp*::pKnock | *M. loti* | ++ | ++ | ++ |
| **SRmD726** | Δ*pfp*::$Nm^R$ *fbp*::pKnock | *B. japonicum* | + | ++ | ++ |
| **SRmD726** | Δ*pfp*::$Nm^R$ *fbp*::pKnock | *L. pneumophila* | ++ | ++ | ++ |
| **SRmD726** | Δ*pfp*::$Nm^R$ *fbp*::pKnock | *A. brasilience* | ++ | ++ | ++ |
| **SRmD726** | Δ*pfp*::$Nm^R$ *fbp*::pKnock | *R. etli* | + | ++ | ++ |
| **SRmD726** | Δ*pfp*::$Nm^R$ *fbp*::pKnock | *D. acidovorans* | ++ | ++ | ++ |
| **SRmD726** | Δ*pfp*::$Nm^R$ *fbp*::pKnock | *A. olearius* | ++ | ++ | ++ |

* Phenotypes are as follows: ++, wild type growth;  -, no growth.
† No *in trans* expression of a *pfp*.

**Table S4.** Bacterial strains and plasmids

| **Strain** | **Genotype and relevant phenotype** | **Reference** |
|---|---|---|
| *Sinorhizobium meliloti* | | |
| Rm1021 | SU47 *str-21* Sm$^R$ | Meade et al. (33) |
| SRmD673 | Rm1021, Δ*pfp* Sm$^R$ | This work |
| SRmD674 | Rm1021, Δ*pfp::*Nm, Sm$^R$ Nm$^R$ | This work |
| SRmD726 | SRmD674, *SMc00535*::pKNOCK Gm, Sm$^R$ Nm$^R$ Gm$^R$ | This work |
| SRmD727 | Rm1021, *SMc00535*::pKNOCK Gm, Sm$^R$ Gm$^R$ | This work |
| *Escherichia coli* | | |
| MM294A | *pro-82 thi-1 hsdR17 supE44* | Finan et al. (3) |
| MT607 | MM294A *recA56* | Finan et al. (3) |
| MT616 | MT607(pRK600) | Finan et al. (3) |
| DH5α | λ$^-$ *ϕ*80d*lacZ*ΔM15 Δ(*lacZYA-argF*)*U169 recA1 endA1 hsdR17*($r_k^-$ $m_k^-$) *supE44 thi-1 gyrA relA1* | Hanahan (3) |
| DH5αRif | Rifampin-resistant variant of DH5α | House et al. (4) |
| DH5αλpir | λ*pir* lysogen of DH5α | House et al. (4) |
| JM109 | *end*A1, *rec*A1, *gyr*A96, *thi, hsd*R17 ($r_k^-$, $m_k^+$), *rel*A1, *sup*E44, Δ(*lac-pro*AB), [F′ *tra*D36, *pro*AB, *laq*I$^q$ZΔM15] | Hanahan (34) |
| **Plasmids** | **Description** | **Reference** |
| pRK600 | pRK2013 *npt*::Tn*9,* Cm$^R$ | Finan et al. (3) |
| pRK7813 | Broad-host-range cloning vector, Tc$^R$ | Jones and Gutterson (35) |
| pCO37 | pRK7813 containing *attB* sites; Gateway-compatible destination vector | Jacob et al. (5) |
| pMK2014 | FRT-*ccdB*-Cm$^r$-FRT cassette, Pen$^R$ | House et al. (4) |
| pXINT129 | λ*int* and *xis* driven by $P_{lac,}$ Km$^R$ | Platt et al. (36) |
| pJQ200SK | Gene replacement suicide vector, Gm$^R$ | Quandt & Hynes (7) |
| pDK36 | pJQ200SK/*pfp* flanking regions, Gm$^R$ | This work |
| pSK11 | pJQ200SK/*pfp* flanking regions with Nm marker, Nm$^R$ Gm$^R$ | This work |
| pRP7 | pKNOCK Gm/*SMc00535* 400bp internal region, Gm$^R$ | This work |
| pMK66 | pCO37/*pfp* from *S. meliloti,* Tc$^R$ | This work |
| pRP20 | pCO37/*SMc00535* Tc$^R$ | This work |
| pRP14 | pUC57/*pfp* from *R. etli*, Amp$^R$ | This work |
| pRP15 | pUC57/*pfp* from *L. pneumophila*, Amp$^R$ | This work |
| pRP16 | pUC57/*pfp* from *B. japonicum*, Amp$^R$ | This work |
| pRP17 | pUC57/*pfp* from *M. loti*, Amp$^R$ | This work |
| pRP18 | pUC57/*pfp* from *A. brasilience*, Amp$^R$ | This work |
| pRP19 | pUC57/*pfkA* from *E. coli*, Amp$^R$ | This work |
| pRP27 | pUC57/*pfp* from *A. olearius*, Amp$^R$ | This work |
| pRP28 | pUC57/*pfp* from *D. acidovorans*, Amp$^R$ | This work |
| pRP21 | PRK7813 containing *pfp* from *M. loti,* Tc$^R$ | This work |
| pRP22 | pRK7813/*pfp* from *B. japonicum,* Tc$^R$ | This work |
| pRP23 | pRK7813/*pfkA* from *E. coli,* Tc$^R$ | This work |

| | | |
|---|---|---|
| pRP24 | pRK7813/*pfp* from *L. pneumophila,* $Tc^R$ | This work |
| pRP25 | pRK7813/*pfp* from *A. brasilience,* $Tc^R$ | This work |
| pRP30 | pRK7813/*pfp* from *D. acidovorans,* $Tc^R$ | This work |
| PRP31 | pRK7813/*pfp* from *A. olearius,* $Tc^R$ | This work |
| pRP32 | pRK7813/*pfp* from *R. etli,* $Tc^R$ | This work |
| pSK16 | pQE9/*pfp* codon optimized for overexpression, $Amp^R$ | This work |
| pRP35 | pQE30/*SMc00535* codon optimized for overexpression, $Amp^R$ | This work |

ΦM12 transducing lysates are denoted by Φ before the strain number. Arrows indicate ΦM12-mediated transduction from the lysate into the recipient strain.
$Sm^R$, streptomycin resistant; $Nm^R$, neomycin resistant; $Gm^R$, gentamicin resistant; $Sp^R$, spectinomycin resistant; $Cm^R$, chloramphenicol resistant; $Tc^R$, tetracycline resistant; $Pen^R$, penicillin resistant; FRT, flippase recognition target.

**Table S5.** Primer sequences

| **Primer name** | **Primer sequence 5’-3’** |
|---|---|
| **Δ*pfp* *** | |
| Left_pfp_um_F | tcctgcagcccgggggatcATTGTAAAGCCTGGCGACG |
| Left_pfp_um_R | gtcgttcaGGGGTCCTCCTGGGGATTTC |
| Right_pfp_um_F | aggaccccTGAACGACGAGGAACGGTTC |
| Right_pfp_um_F | gaacaaaagctggagctcATGCCGAGCCAGACGAGATAG |
| **Δ*pfp*-Nm** | |
| pfp_L_BamHI_F | cgaattcctgcagcccggggCCGTAAGCGCGTTACCAG |
| pfp_L_BamHI_R | tatggctcatGGGGTCCTCCTGGGGATTTC |
| pfp_R_SacI_F | aatttttctaaTGAACGACGAGGAACGGTTC |
| pfp_R_SacI_R | agggaacaaaagctggagctTCTCGAAGATCGCCATCTG |
| Neo_BamSac_F | ggaggaccccATGAGCCATATTCAGCGTG |
| Neo_BamSac_R | tcctcgtcgttcaTTAGAAAAATTCATCCAGCATC |
| del_pfp_confir_F | TTCCTGGACGCCACCGTGC |
| del_pfp_confir_R | GAATGGCAAGCAGCACCGCA |
| ***smc00535*::pKNOCK GmR** | |
| 535int_PstI_F | AACTGCAGGTTCATCGGCGAGGAATCCG |
| 535int_BamHI_R | TTGGATCCCGCCGCGCAGCGATA |
| 535Lflank_FWD | ATTCGCCGCGGCCATGGAGT |
| pKnockMCS_FWD | AACCCTCATGGCTAACGTACTAAGCT |
| **Codon optimized *pfp*** | |
| Pfp_Bam_FW | GATCGGATCCGCAAAAAGAAAGTAGCTAT |
| Pfp_Hind_RV | GATCAAGCTTTTAGTCAGCGGTCTGCCACT |
| **Insertion-sequencing †** | |
| 1TN-mariner | GTTCGCTTGCTGTCC |
| 1GG | CAGACGTGTGCTCTTCCGATCggggggggggg |
| 2TNA-mariner | AATGATACGGCGACCACCGAGATCTACACTCTTTCCCTACA<br>CGACGCTCTTCCGATCTcggggacttatcatccaacc |
| 2TNB-mariner | AATGATACGGCGACCACCGAGATCTACACTCTTTCCCTACA<br>CGACGCTCTTCCGATCTAcggggacttatcatccaacc |
| 2TNC-mariner | AATGATACGGCGACCACCGAGATCTACACTCTTTCCCTACA<br>CGACGCTCTTCCGATCTGAcggggacttatcatccaacc |
| 2BAR5 | CAAGCAGAAGACGGCATACGAGATCACTGTGTGACTGGAGT<br>TCAGACGTGTGCTCTTCCGATC |
| 2BAR6 | CAAGCAGAAGACGGCATACGAGATGATCTGGTGACTGGAGT<br>TCAGACGTGTGCTCTTCCGATC |
| 2BAR7 | CAAGCAGAAGACGGCATACGAGATTGGTCAGTGACTGGAGT<br>TCAGACGTGTGCTCTTCCGATC |
| 2BAR8 | CAAGCAGAAGACGGCATACGAGATCTGATCGTGACTGGAGT<br>TCAGACGTGTGCTCTTCCGATC |

* Lower case letters represent homologous overlap for Gibson assembly.

† Lower case letters represent regions of homology to the template, while the underlined regions represent an Illumina index sequence.

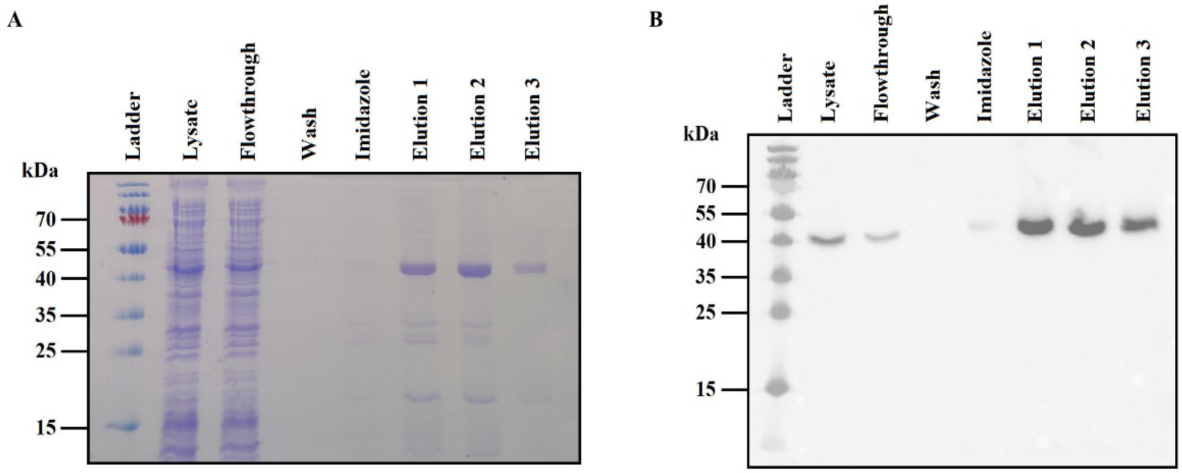


**Figure S1**. **IMAC purification of overexpressed His-tagged Pfp.** (**A**) Elutants separated and visualized by SDS-PAGE stained with Coomassie Brilliant Blue. The PageRuler$^{TM}$ pre-stained protein ladder was included to distinguish Pfp by molecular weight (kDa) and is shown on the left. (**B**) Western blot analysis of SDS-PAGE with RGS-His HRP conjugate to detect Pfp in the elutant samples.

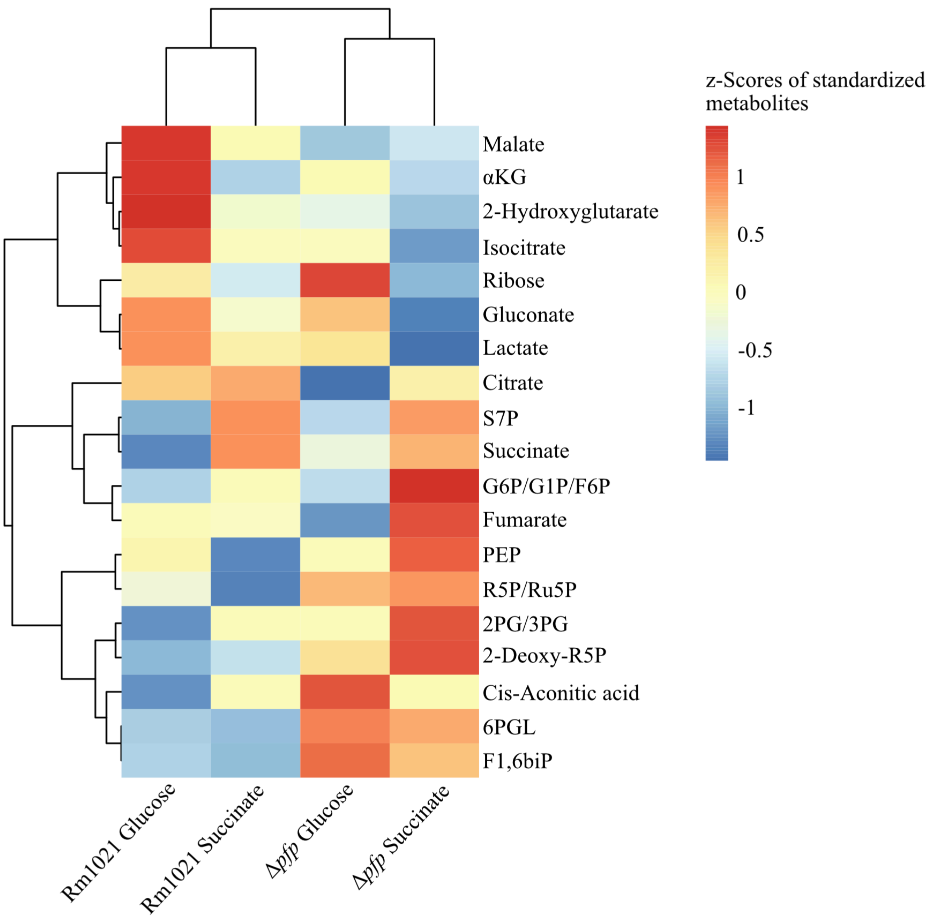


**Figure S2. Impact of deleting *pfp* on metabolite concentrations in *Sinorhizobium meliloti*.** A hierarchical clustered heat map of central carbon metabolites of *S. meliloti*. Columns are representative of wildtype *S. meliloti* Rm1021 or a *S. meliloti* Δ*pfp* mutant (strain SRmD674) grown with glucose or succinate as the sole carbon source. Colour gradients across each row correspond to the fold change with respect to the z-score for the mean of each metabolite pool. 2-phosphoglycerate (2-PG); 3-phosphoglycerate (3-PG); phosphoenolpyruvate (PEP); glucose-6-phosphate (G6P); glucose-1-phosphate (G1P); fructose-6-phosphate (F6P); α-ketoglutarate (αKG); ribose-5-phosphate (R5P); ribulose-5-phosphate (Ribu5P); Sedoheptulose-7-phosphate (S7P); fructose-1,6-bisphosphate (F1,6biP); 6-phosphogluconate. (6PG).

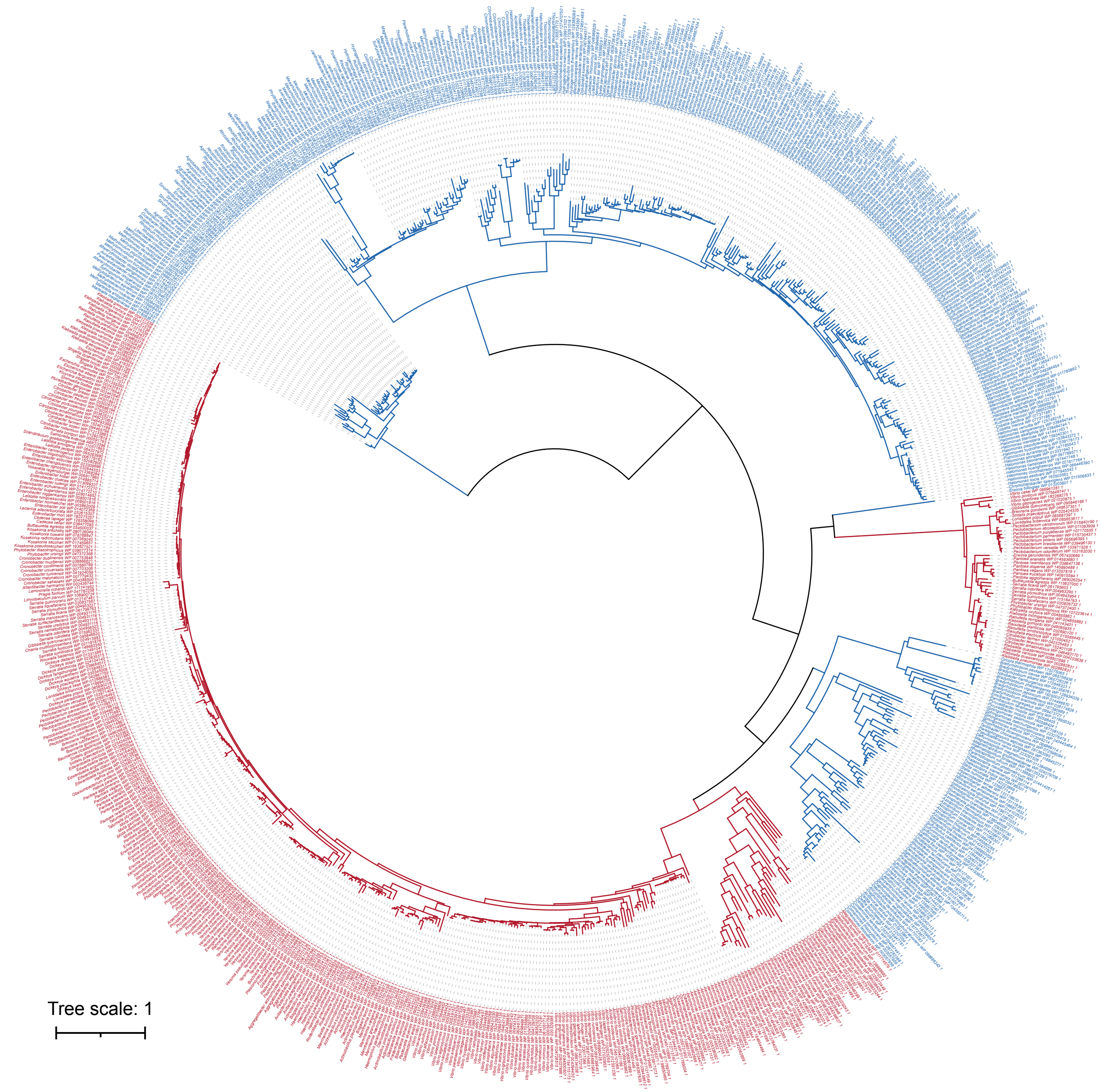


**Figure S3. A maximum likelihood phylogeny of putative phosphofructokinases.** An unrooted, maximum likelihood phylogeny of the 800 putative phosphofructokinase enzymes identified in this study is shown. The scale bar represents the average number of amino acid substitutions per site. Clades are colour coded based on whether the enzyme is predicted to use ATP (red) or pyrophosphate (blue) as a substrate. An interactive version of this phylogeny, with support values, can be accessed at itol.embl.de/shared/Xjwm5Fs4ugTK.

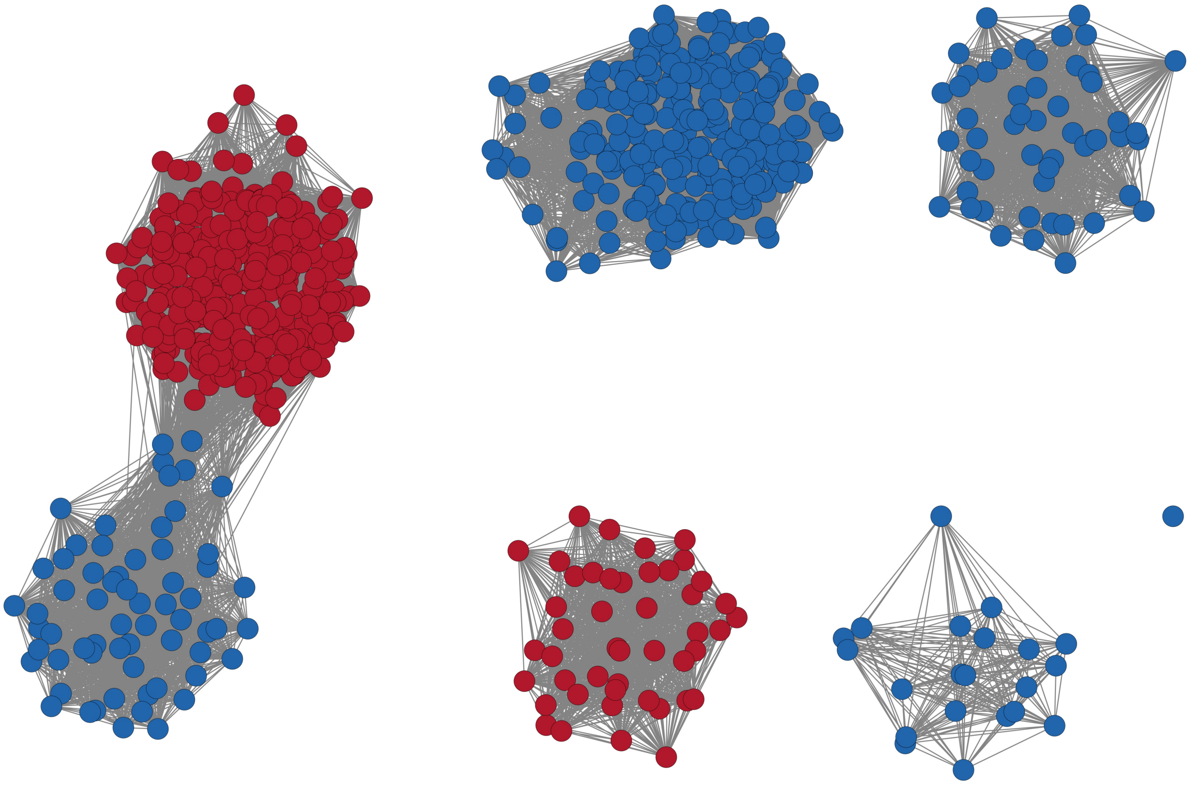

**Figure S4. A sequence similarity network of putative phosphofructokinases.** A sequence similarity network, calculated using EFI-EST, of the 800 putative phosphofructokinase enzymes identified in this study is shown. Each node (the circles) represents one protein, while edges (the lines) indicate that the sequence similarity between pairs of proteins exceeds the threshold. Nodes are colour coded based on whether the enzyme is predicted to use ATP (red) or pyrophosphate (blue) as a substrate.

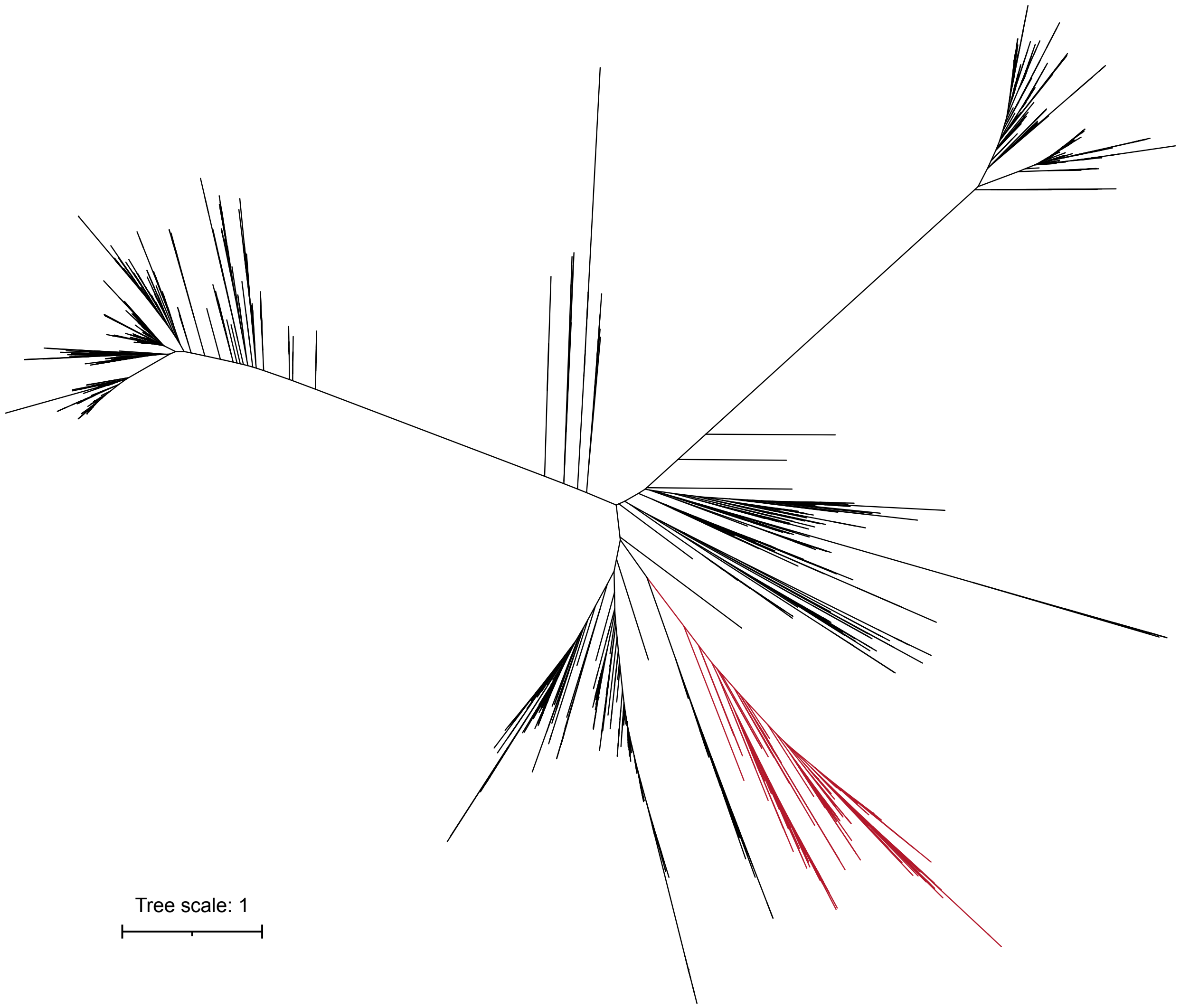


**Figure S5. A maximum likelihood phylogeny of putative fructose-1,6-bisphosphatases.** An unrooted, maximum likelihood phylogeny of the 2,810 putative fructose-1,6-bisphosphatase enzymes identified in this study is shown. The scale bar represents the average number of amino acid substitutions per site. The clade containing Smc00535 is coloured red. An interactive version of this phylogeny, with support values, can be accessed at itol.embl.de/shared/Xjwm5Fs4ugTK.

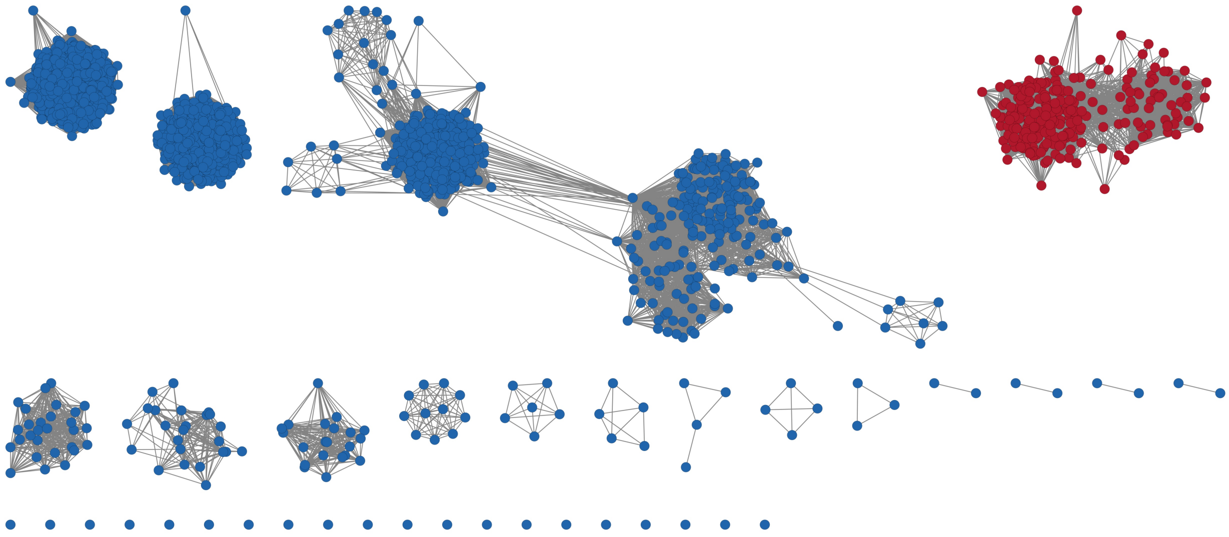

**Figure S6. A sequence similarity network of putative fructose-1,6-bisphosphatases.** A sequence similarity network, calculated using EFI-EST, of the 2,810 putative phosphofructokinase enzymes identified in this study is shown. Each node (the circles) represents one protein, while edges (the lines) indicate that the sequence similarity between pairs of proteins exceeds the threshold. Nodes of the cluster containing Smc00535 are coloured red; all other nodes are in blue.

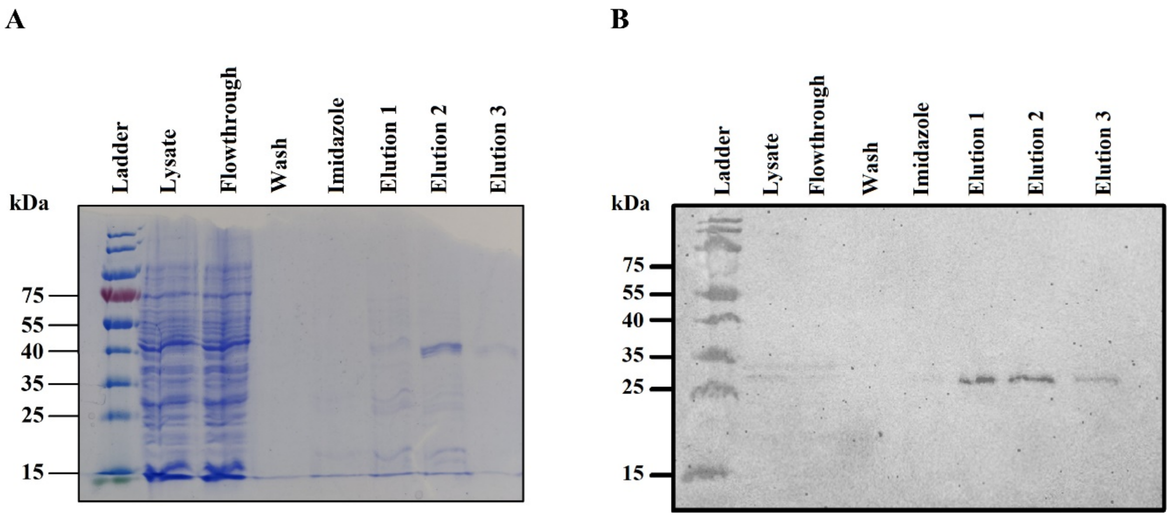


**Figure S7. Verification of IMAC purification of overexpressed His-tagged Smc00535 (Fbp).** (**A**) Elutants separated by SDS-PAGE stained with Coomassie Brilliant Blue. The PageRuler$^{TM}$ pre-stained protein ladder was included to distinguish Fbp by molecular weight (kDa) and is shown on the left. (**B**) Western blot analysis of SDS-PAGE with RGS-His HRP conjugate to detect Fbp in the elutants.

### SUPPLEMENTARY DATASETS

**Dataset S1. Quantification of intracellular metabolites via LC-MS.** Rows contain data corresponding to a standard or sample. Columns A-H contain user-defined settings for LS-MS and assayed samples. 25x denoted in data file names indicate a 25-fold sample dilution prior to analysis. Subsequent columns are grouped by the metabolite listed in row 1 and list LS-MS measurement readings for each row. Cells are colour-coded based on the quality of quantification indicated below the recorded data. The notes sheet provides information regarding suggested dilution calculations. Overall comments regarding the quality of each metabolite run are listed.

**Dataset S2. The complete Insertion-sequencing dataset.** Each row contains data for one gene. Columns are defined as follows: Locus_tag – the gene's locus tag in the GenBank annotation; Gene_name – the name of the gene, where available, or otherwise the locus tag; Replicon_name – the name of the replicon encoding the gene (chromosome, pSymA, pSymB); Replicon_accession – the replicon's GenBank accession number; Strand – whether the gene is on the plus or minus strand; Gene_start – the nucleotide start position for the gene; Gene_end – the nucleotide end position for the gene; TA_site_total – the total number of TA sites within the central 90% of a gene, regardless of whether it contained insertions; TA_site_hits – the number of TA sites within the central 90% of a gene that had at least one mapped read in the given sample; Read_count – the total number of reads mapping to TA sites in the central 90% of a gene in the given sample; Normalized_read_count – the number of reads mapping to TA sites in the central 90% of a gene in the given sample, normalized by sequencing depth; Median_normalized_hit_count – median number of reads per TA site with at least one mapped read per gene, normalized by sequencing depth; Summary_score – a statistic calculated by multiplying the median normalized hit count by the number of TA sites in the gene with at least one mapped read, divided by the total number of TA sites in the central 90% of the gene; WT-Glucose – data for *S. meliloti* RmP110 grown with glucose as the carbon source; WT-Succinate - data for *S. meliloti* RmP110 grown with succinate as the carbon source; *pfp*-Glucose - data for *S. meliloti* Δ*pfp* grown with glucose as the carbon source; *pfp*-Succinate - data for *S. meliloti* Δ*pfp* grown with succinate as the carbon source.

**Dataset S3. Genomes used in this study.** A lists of the 1,408 genomes used to infer the species phylogeny, and whose proteomes were searched for phosphofructokinase and fructose-1,6-bisphosphatase proteins. Columns are defined as follows: species – the genus and species of the organism; strain – the strain name of the organism; class – the taxonomic classification of the organism at the class level; phosphofructokinase – indicates if the organism is predicted to encode an ATP-dependent phosphofructokinase, a pyrophosphate (PPi)-dependent phosphofructokinase, both types of enzymes, or neither; Smc00535_ortholog – indicates if the organism is predicted to encode at least one Smc00535 ortholog (Yes) or not (No); NCBI_Assembly_accession – the NCBI Assembly accession for the genome; Ftp_link – a ftp link to download the genome from NCBI.

**Dataset S4. Putative phosphofructokinase enzymes detected in this study.** A list of the 800 putative phosphofructokinases identified in this study. Columns are defined as follows: species – the genus and species name of the organism encoding the phosphofructokinase; NCBI_Protein_accession – the NCBI Protein accession number for the protein; Predicted_substrate_specificity – Whether the enzyme is predicted to use ATP or pyrophosphate (PPi) as a substrate.

**Dataset S5. Putative fructose-1,6-bisphosphatase enzymes detected in this study.** This file lists the 2,810 putative fructose-1,6-bisphosphatases identified in this study. Columns are defined as follows: species – the genus and species name of the organism encoding the phosphofructokinase; NCBI_Protein_accession – the NCBI Protein accession number for the protein; Predicted_Smc00535_ortholog? – whether the protein is predicted to be an ortholog of Smc00535 (Yes) or not (No).

### SI REFERENCES

1. J. Sambrook, E. F. Fritsch, T. Maniatis, *Molecular cloning: A laboratory manual* (Cold Spring Harbor Laboratory, 1989).

2. T. M. Finan, *et al.*, General transduction in *Rhizobium meliloti*. *J. Bacteriol.* **159**, 120–124 (1984).

3. T. M. Finan, B. Kunkel, G. F. De Vos, E. R. Signer, Second symbiotic megaplasmid in *Rhizobium meliloti* carrying exopolysaccharide and thiamine synthesis genes. *J. Bacteriol.* **167**, 66–72 (1986).

4. B. L. House, M. W. Mortimer, M. L. Kahn, New recombination methods for *Sinorhizobium meliloti* genetics. *Appl. Environ.l Microbiol.***70**, 2806–2815 (2004).

5. A. I. Jacob, *et al.*, Mutational analysis of the *Sinorhizobium meliloti* short-chain dehydrogenase/reductase family reveals substantial contribution to symbiosis and catabolic diversity. *MPMI* **21**, 979–987 (2008).

6. M. F. Alexeyev, The pKNOCK series of broad-host-range mobilizable suicide vectors for gene knockout and targeted DNA insertion into the chromosome of gram-negative bacteria. *BioTechniques* **26**, 824–828 (1999).

7. J. Quandt, M. F. Hynes, Versatile suicide vectors which allow direct selection for gene replacement in gram-negative bacteria. *Gene* **127**, 15–21 (1993).

8. M. M. Bradford, A rapid and sensitive method for the quantitation of microgram quantities of protein utilizing the principle of protein-dye binding. *Anal. Biochem.* **72**, 248–254 (1976).

9. U. K. Laemmli, Cleavage of structural proteins during the assembly of the head of bacteriophage T4. *Nature* **227**, 680–685 (1970).

10. H. Towbin, T. Staehelin, J. Gordon, Electrophoretic transfer of proteins from polyacrylamide gels to nitrocellulose sheets: procedure and some applications. *Proc. Natl. Acad. Sci. U.S.A.* **76**, 4350–4354 (1979).

11. B. J. Perry, C. K. Yost, Construction of a mariner-based transposon vector for use in insertion sequence mutagenesis in selected members of the *Rhizobiaceae*. *BMC Microbiol.* **14**, 298 (2014).

12. G. C. diCenzo, *et al.*, Robustness encoded across essential and accessory replicons of the ecologically versatile bacterium *Sinorhizobium meliloti*. *PLoS Genet.* **14**, undefined-undefined (2018).

13. P. A. Cowie, *et al.*, Investigating the surface process response to fault interaction and linkage using a numerical modelling approach. *Basin Res.* **18**, 231–266 (2006).

14. B. Bushnell, BBMap: A Fast, Accurate, Splice-Aware Aligner. (2014).

15. A. M. Bolger, M. Lohse, B. Usadel, Trimmomatic: a flexible trimmer for Illumina sequence data. *Bioinformatics* **30**, 2114–2120 (2014).

16. F. Galibert, *et al.*, The composite genome of the legume symbiont *Sinorhizobium meliloti*. *Science* **293**, 668–672 (2001).

17. B. Langmead, S. L. Salzberg, Fast gapped-read alignment with Bowtie 2. *Nat. Methods* **9**, 357–359 (2012).

18. P. Danecek, *et al.*, Twelve years of SAMtools and BCFtools. *Gigascience* **10**, giab008 (2021).

19. G. C. Dicenzo, *et al.*, Multidisciplinary approaches for studying rhizobium–legume symbioses. *Can. J. Microbiol.* **65**, 1–33 (2019).

20. M. Wu, A. J. Scott, Phylogenomic analysis of bacterial and archaeal sequences with AMPHORA2. *Bioinformatics* **28**, 1033–1034 (2012).

21. K. Katoh, D. M. Standley, MAFFT multiple sequence alignment software version 7: improvements in performance and usability. *Mol. Biol. Evol.* **30**, 772–780 (2013).

22. S. Capella-Gutiérrez, J. M. Silla-Martínez, T. Gabaldón, trimAl: a tool for automated alignment trimming in large-scale phylogenetic analyses. *Bioinformatics* **25**, 1972–1973 (2009).

23. S. Kalyaanamoorthy, B. Q. Minh, T. K. F. Wong, A. von Haeseler, L. S. Jermiin, ModelFinder: fast model selection for accurate phylogenetic estimates. *Nat. Methods* **14**, 587–589 (2017).

24. L.-T. Nguyen, H. A. Schmidt, A. von Haeseler, B. Q. Minh, IQ-TREE: A fast and effective stochastic algorithm for estimating maximum-likelihood phylogenies. *Mol. Biol. Evol.* **32**, 268–274 (2015).

25. S. Guindon, *et al.*, New algorithms and methods to estimate maximum-likelihood phylogenies: assessing the performance of PhyML 3.0. *Syst. Biol.* **59**, 307–321 (2010).

26. I. Letunic, P. Bork, Interactive tree of life (iTOL) v3: an online tool for the display and annotation of phylogenetic and other trees. *Nucleic Acids Res.* **44**, W242-245 (2016).

27. R. D. Finn, *et al.*, The Pfam protein families database: towards a more sustainable future. *Nucleic Acids Res.* **44**, D279–D285 (2016).

28. D. H. Haft, *et al.*, TIGRFAMs and genome properties in 2013. *Nucleic Acids Res*. **41**, D387-395 (2013).

29. S. R. Eddy, A new generation of homology search tools based on probabilistic inference. *Genome. Inform.* **23**, 205–211 (2009).

30. R. Zallot, N. Oberg, J. A. Gerlt, The EFI web resource for genomic enzymology tools: leveraging protein, genome, and metagenome databases to discover novel enzymes and metabolic pathways. *Biochemistry* **58**, 4169–4182 (2019).

31. N. Oberg, R. Zallot, J. A. Gerlt, EFI-EST, EFI-GNT, and EFI-CGFP: Enzyme function initiative (EFI) web resource for genomic enzymology tools. *J. Mol. Biol.* **435**, 168018 (2023).

32. P. Shannon, *et al.*, Cytoscape: a software environment for integrated models of biomolecular interaction networks. *Genome Res*. **13**, 2498–2504 (2003).

33. H. M. Meade, S. R. Long, G. B. Ruvkun, S. E. Brown, F. M. Ausubel, Physical and genetic characterization of symbiotic and auxotrophic mutants of *Rhizobium meliloti* induced by transposon Tn5 mutagenesis. *J. Bacteriol.* **149**, 114–122 (1982).

34. D. Hanahan, Studies on transformation of *Escherichia coli* with plasmids. *J. Mol. Biol.* **166**, 557–580 (1983).

35. J. D. Jones, N. Gutterson, An efficient mobilizable cosmid vector, pRK7813, and its use in a rapid method for marker exchange in *Pseudomonas fluorescens* strain HV37a. *Gene* **61**, 299–306 (1987).

36. R. Platt, C. Drescher, S.-K. Park, G. J. Phillips, Genetic system for reversible integration of DNA constructs and *lacZ* gene fusions into the *Escherichia coli* chromosome. *Plasmid* **43**, 12–23 (2000).
